# New Insights Into the Effect of Residue Mutation on the Rotavirus VP1 Function Using Molecular Dynamic Simulations

**DOI:** 10.1101/2020.04.08.031443

**Authors:** Nabil Abid, Marco Salemi, Giovanni Chillemi

**Affiliations:** Laboratory of Transmissible Diseases and Biological Active Substances LR99ES27, Faculty of Pharmacy, University of Monastir, Rue Ibn Sina, 5000, Monastir, Tunisia; High Institute of Biotechnology of Sidi Thabet, Department of Biotechnology, University Manouba, BP-66, 2020, Ariana-Tunis, Tunisia; Department of Pathology, Immunology and Laboratory Medicine, University of Florida College of Medicine, Emerging Pathogens Institute, Gainesville, University of Florida, P.O. Box 100009, FL 32610-3633, USA; Department for Innovation in Biological, Agro-food and Forest systems, DIBAF, University of Tuscia, via S. Camillo de Lellis s.n.c., 01100 Viterbo, Italy; Institute of Biomembranes, Bioenergetics and Molecular Biotechnologies, IBIOM, CNR, Via Giovanni Amendola, 122/O, 70126, Bari, Italy

**Keywords:** Rotavirus RdRp, Molecular dynamics simulation, Bioinformatics, Mutations

## Abstract

Rotavirus group A remains a major cause of diarrhea in infants and young children worldwide. The permanently emergence of new genotypes puts the potential effectiveness of vaccines under serious question. Thirteen VP1 mutants were analyzed using molecular dynamic simulations and the results were combined with the experimental findings, reported previously. The results revealed structural fluctuations and secondary structure change of VP1 protein that may alter its function during viral replication/transcription. Altogether, the structural analysis of VP1 may boost efforts to develop antivirals, as they might complement the available vaccines.

## 1. Introduction

Rotavirus A (RVA) is a double-stranded RNA (dsRNA) virus of the *Reoviridae* family. It is a significant cause of childhood gastroenteritis and accounts for ∼450,000 deaths annually, most occurring in developing countries [1]. RVA causes also great economic loss to livestock industry worldwide, and again mostly in the developing countries [2,3]. The structural analysis of RVA proteins upon mutations may enhance our ability to better understand their role in viral replication. We focus in the present study on rotavirus RNA-directed RNA polymerase (RdRp), named VP1 protein and coded by VP1 gene, as one of the most important proteins in the viral replication; it is involved in both transcription and genome replication/packaging [4,5] in the presence of other proteins, such as VP2 protein; its N-terminal tether domain extends around VP1 and forms cradle that helps stabilize the enzyme in position [6], suggesting to regulate VP1 transitions between transcription and replication states [7].

From the structural point of view, VP1 protein is organized as three distinct domains: an N-terminal domain (residues 1 to 332); a central polymerase domain with canonical fingers (residues 333 to 488 and residues 524 to 595), palm (residues 489 to 523 and residues 596 to 685), and thumb subdomains (residues 686 to 778); and a C-terminal “bracelet” domain (residues 779 to 1089) [8]. Together, the N- and C-terminal domains enclose the central polymerase domain to create a cage-like enzyme with a buried active site. Although the crystal structures of the RVA VP1 have been determined and fitted into their capsid structures [7-9], the structures of the viral particle-associated VP1 and cofactor proteins, such as NSP2, NSP5 and VP3, are still unknown.

The importance of amino acid residues that affect the activity of VP1 protein has been demonstrated [7,10-12] and mutant VP1 proteins in specific positions were engineered in order to assess their capacity to synthesize dsRNA *in vitro* [10,11]. Authors reported that, based on their effect on the replication level, the induced mutations can be classified into three groups: (i) mutations with no significant effect on replication level; (ii) mutations significantly lower levels of dsRNA product; and (iii) mutation that enhanced the initiation capacity and product elongation rate of VP1 protein. In addition, temperature-sensitive (ts) mutant mapped to the gene encoding VP1 protein was generated via chemical mutagenesis of prototypic strain SA11 [13]. This mutant shows diminished viral growth, RNA synthesis, protein synthesis, and virion morphogenesis at 39°C compared to that at 31°C [13-15].

The three dimensional structure of proteins is not static, the different residues can move, and thus, stabilize or break non-covalent interactions. To better understand the effect of the generated mutants, structural data need to be linked to the temporal context of the virus replication/transcription due to the presence of a recently published structure for a catalytically active form of VP1 protein [7,9].

We sought to determine how VP1 residues mutations at the RNA entry tunnel contribute to the decrease of the dsRNA synthesis, based on previous experimental findings. Toward this end, molecular dynamics (MD) simulation were used to simulate protein movements at atomic resolution. The resulted genomic and structural data were analyzed and combined with the new available structural and experimental data to investigate the effects of these mutations on the VP1 protein structure, dynamics and on its ability to function properly.

## 2. Materials and Methods

### 2.1. Structure preparation

Experimentally determined structures of strain SA11 VP1 [PDB ID: 2R7R] was retrieved from RCSB Protein Data Bank (https://www.rcsb.org/) for our study. To create VP1 for simulations, the nucleic acid (chain X) was removed from the input structure and the missing flexible loop (residues 346 to 358) was modeled with the Modeler loop-modeling tool embedded in the program UCSF Chimera v1.12 [16] using locally installed Modeller software [17] using the loop modeling protocol of Discrete Optimized Protein Energy (DOPE) [18]. Out of the 10 generated models, the model with the most favorable zDOPE score was chosen for further computations.

### 2.2. VP1 Mutant generation and molecular dynamics simulation

The selected residues at the RNA entry tunnel and the catalytic site were shown in (**Table 1, Video S1**). To create mutant VP1 structures for simulations, Chimera’s Rotamers tool was used to replace the native residue in the VP1 structure [19]. The residue rotamer from the Dunbrack backbone-dependent rotamer library with the highest probability (if required) was chosen for each mutant VP1 model.

**Table 1.**
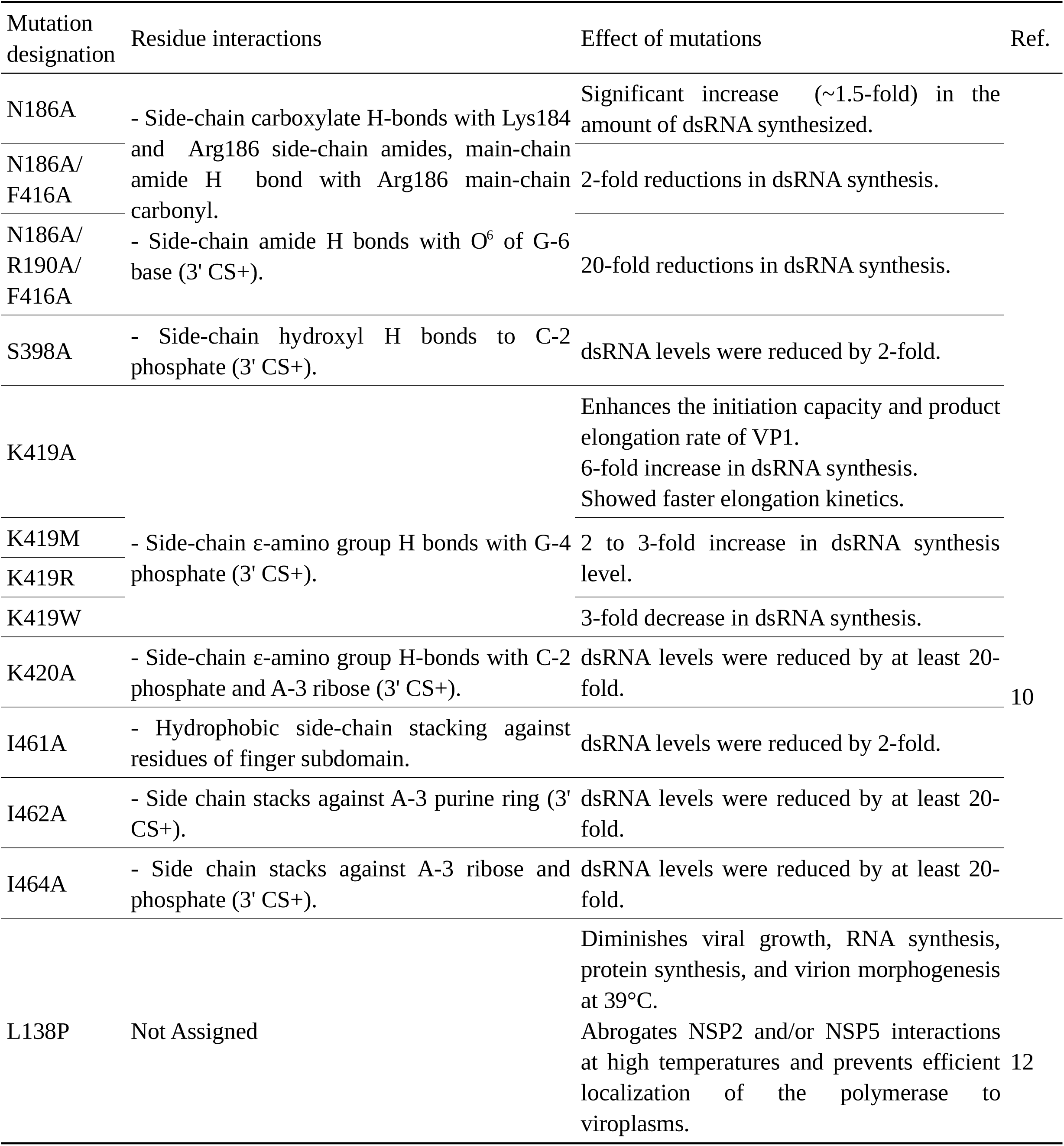
The list of mutants used in the present study.

The starting structures were embedded in a triclinic box, extending up to 15 Å from the solute, and immersed in SPC water molecules [20]. Counter ions were added to neutralize the overall charge with the genion gromacs tool. After energy minimizations, the systems were slowly relaxed for 5 ns by applying positional restraints of 1000 kJ mol^-1^ nm^-2^ to the protein atoms. Then unrestrained MD simulations were carried out for a length of 225 ns with a time step of 2 fs with GROMACS 2018.3 simulation package [21] (supercomputer Galileo, CINECA, Bologna, Italy) and the gromos54a7.ff force field [22]. V-rescale temperature coupling was employed to keep the temperature constant at 300 K [23]. The Particle-Mesh Ewald method was used for the treatment of the long-range electrostatic interactions [24]. The first 5 ns portion of the trajectory was excluded from the analysis.

### 2.3. MD trajectory analysis of VP1 structures

The simulation trajectories were analyzed using several auxiliary programs provided with the GROMACS 2018.3 package. These programs include *gmx rmsd, gmx rmsf, gmx hbond, gmx gyrate, gmx sasa*, and *gmx do_dssp* utilities for the analysis of root mean square deviations (RMSDs), root mean square fluctuations (RMSFs), hydrogen bonds (H-bonds), radius of gyration (R_gyr_), solvent-accessible surface area (SASA), and secondary structure profile (SS), respectively. Distance between residues was performed using distance script in VMD program [25].

The convergence of the MD trajectories was assessed by calculation of RMSD from the starting structures as a function of simulation time. The RMSD value, a measure of molecular mobility, is calculated by translating and rotating the coordinates of the instantaneous structure to superimpose the reference with a maximum overlap. The RMSD is mathematically defined as:

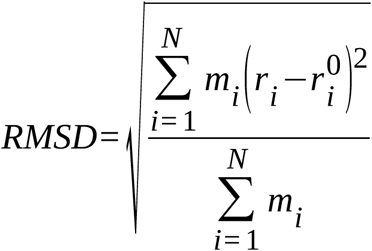

where m_i_ is the mass of atom i. *r*_*i*_ and 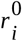 are the coordinates of atom i at a certain instance during MD simulations and at its reference state, respectively.

The conformational flexibility of both proteins during their corresponding MD trajectories was assessed by means of RMSF calculations. RMSF values were calculated after superimposing each individual structure of a trajectory onto the initial structure by means of least-squares fitting, to remove rotational and translational motions. The RMSF values are a measure of atomic fluctuations along an MD trajectory and were calculated using the equation:

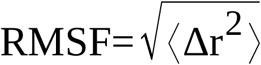

where Δr is the instantaneous fluctuation (displacement) of the position of an atom (or set of atoms) relative to its mean position and <> denotes simulation time averaged values over the MD trajectory. RMSF values were calculated after superimposing each individual structure of a trajectory onto the initial structure by means of least-squares fitting, to remove rotational and translational motions.

H-bonds were calculated between hydrogen donors and acceptors, considering to be formed if the donor-to-acceptor distance is fewer than 0.35 nm and the donor-hydrogen-acceptor angle is within 30° of linearity.

The R_gyr_ measures the compactness of a structure. It is defined as the mass-weighted geometric mean of the distance of each atom from the protein’s center of mass. The R_gyr_ was computed using all the structure atoms in the standard formula:

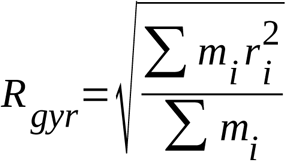

where r_i_ is the distance of atom i from the center of mass of the enzyme, and m_i_ is its mass.

The *gmx do_dssp* utility works by calculating the most likely secondary structure assignment given the 3D structure of a protein. It does this by reading the position of the atoms in a protein followed by calculation of the H-bond energy between all atoms. The algorithm will discard any hydrogen present in the input structure and calculates the optimal hydrogen positions by placing them at 1.00 Å from the backbone N in the opposite direction from the backbone C=O bond. The best two H-bonds for each atom are then used to determine the most likely class of secondary structure for each residue in the protein.

For additional support of our MD simulation results, we collected the large-scale motions of the Native and mutant proteins using essential dynamics (ED) analysis or principal component analysis (PCA).

The PCA method is based on the construction of the covariance matrix of the coordinate fluctuations of the simulated proteins. PCA uses the covariance matrix of atomic coordinates, the elements of which are defined as:

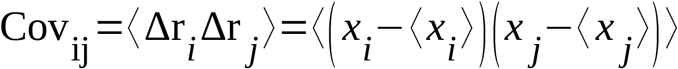

where Δr_i_ and Δr_j_ are the vectors of the instantaneous fluctuation of the position of atoms i and j, relative to their mean positions and <> denotes time averaged values over the MD trajectory. Diagonalization of the covariance matrix produces a set of eigenvectors, each defined by an eigenvalue, corresponding to directions and amplitudes of collective motions. Thus, PCA allows elimination of the noise from the dominant modes of system motions. *xi* and *xj* are atomic coordinates of each Cα atom, and the brackets denote the average. Eigenvectors with the largest eigenvalues are representative of the slowest modes, and generally are associated with large-scale movements in proteins, which are responsible for protein function.

The *gmx-covar* function was used for the construction and diagonalization of Cov_ij_ whereas the *gmx-anaeig* function was used to project the MD trajectories onto the main eigenvectors, maintaining the covariance matrix as a starting point. The PCA scatter plots were generated using these Gromacs built-in utilities [26]. The trajectory files were analyzed, and the graph was plotted using the GRACE Program. Visualization of the atomic model, including figures and movies, is made with Chimera v1.12 [16].

### 2.4. Dynamic cross-correlation analysis

The dynamic cross-correlation (DCC) maps of each system were calculated based on the Cα atoms of residues using *calc_correlation.py* script in the MD-TASK package [27]. Each cell value (C_ij_) in the matrix of the DCC map were calculated using the following formula:

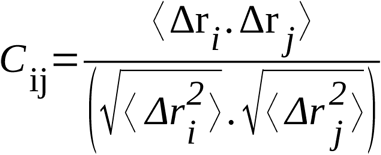

With Δr_i_ represents the displacement from the mean position of atom i, and < > denotes the time average over the whole trajectory. Positive values of C_ij_ show correlated motion between residues i and j, moving in the same direction, whereas negative values of C_ij_ show anti-correlated motion between residues i and j, moving in the opposite direction.

In this study, only Cα was applied for analysis by averaging motions of Cα atoms deviating from the mean structure based on the 225 ns trajectories from MD simulations with a total of 10,000 snapshots.

### 2.5. Network analysis

Networks are an intuitive way of representing relational data using nodes and edges. To analyse the difference in the intra-domain communication upon mutations, the VP1 protein is represented as a residue interaction network (RIN), where the Cβ atoms of each residue (*C*α for Glycine) are treated as nodes within the network, and edges between nodes defined within a distance cut off of 7 Å. In this manner, the RIN was constructed as a symmetric *N* × *N* matrix, where the *ij*^th^ element is assigned as 1 if residue *i* is connected to residue *j* and a zero if no connection exists.

In this study, MD-TASK [27] was used to construct dynamic residue networks (DRN) for each MD trajectory (225 ns), in which RINs are constructed for every *n*th frame of the trajectory using a 10,000 ps time interval, to build a DRN matrix. MD-TASK constructs a DRN and uses this to calculate the changes in betweenness centrality (BC) and average shortest path (*L*) to residues over the trajectory.

#### 2.5.1. Betweenness centrality (BC)

BC is a measure of how important a residue is for communication within a protein. The BC of a node/residue is equal to the number of shortest paths for all nodes to all others that pass through that node [28]. It provides a measure of usage frequency for each node during navigation of the network. BC was calculated using MD-TASK based on Dijkstra’s algorithm [29].

#### 2.5.2. Average shortest path (L)

For a given residue, *L* is the sum of the shortest paths to that residue, divided by the total number of residues less one [30]. By iterating over the DRN, each RIN is analysed in terms of the average of shortest path length (*L*_*ij*_) between residue *i* and any other residue *j.* The average *L*_*ij*_ is then calculated as the average number of steps that the node (residue) may be reached from all other residues in the RIN:

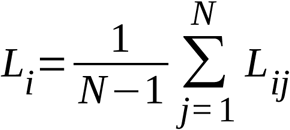

#### 2.5.3. Residue contact map

Residue contact maps are generated by monitoring the interactions of a residue throughout a simulation, yielding a network diagram with the residue of interest [e.g. single nucleotide polymorphism (SNP)] at the center, and residues that it interacts with arranged around it. Edges between the residue of interest and the other residues are weighted based on how often the interaction exists. Inter-domain contacts were evaluated using *contact_map.py* script from MD-TASK with a distance cut-off of 7 Å [27]. The script was utilized every 10,000 ps and the resultant data collated into a single data frame for plotting purposes.

### 2.6. Analysis of Tunnels in MD Trajectories

CAVER Analyst 2.0 [31] was used for the calculation of tunnels in the 10-ns MD simulations of VP1 Native and mutants. The initial starting point of the tunnel search was automatically optimized to prevent its collision with protein atoms. The tunnel search was performed using a probe of 1.5 Å radius and a cost function exponent of 2. Redundant tunnels in each frame were removed using a linear transformation coefficient of 1 and a threshold of 1. Clustering of the tunnels identified during each simulation was performed by hierarchical average link clustering based on the pairwise distances of the tunnels computed using a linear transformation coefficient of 1 and a distance threshold of 3.5. Other parameters used default settings throughout the calculations.

## 3. Results

### 3.1. Analysis of molecular dynamics simulation of VP1 structures

We found that all mutant proteins (**Table 1**) are largely stable and do not induce a globally destabilizing effect in VP1 protein. The greatest portion of RMSDs of all the Cα atoms was reached after approximately 5 ns simulation time, even if further fluctuation in its value are observed in several mutants at later times (**Figure S1**). The R_gyr_ which depicts the compactness of the system, a property linked to the molecular volume and compactness, was shown stable in all the mutant structures (data not shown). All the following analyses were carried out discarding the first 5 ns of simulation time.

We have investigated the changes in secondary structure of Native and mutants. The mutations, except for the 138P mutant, do not induce any significant global change in the secondary structure content and the stable conformation was maintained throughout simulation (Data not shown).

The SASA quantifies the amount of exposure of a given region to the solvent medium. The question whether MD simulations cause different SASA patterns in the mutant structures than the Native structure was assessed. The results showed that the mean SASA values are almost identical (data not shown).

#### 3.1.1. RNA entry bottleneck moderates RNA replication through Lys419

Lys419 is located at the narrowest region of the template entry tunnel, a site that immediately precedes the hollow catalytic center of VP1 protein. It was shown that K419A, K419M, and K419R mutants enhanced capacity to form initiation complexes and dsRNA synthesis; whereas the K419W decreased the dsRNA synthesis by at least 3-fold [10].

Changes in root mean square fluctuations (ΔrRMSF) of backbone atoms in K419A, K419M, K419R, and K419W were analyzed. Positive values of ΔrRMSF correspond to the flexible zone of K419W as compared with K419A, K419M, or K419R; while negative values of ΔrRMSF corresponds to a rigid zone of K419W, also when compared to K419A, K419M, or K419R. The positive values were represented in **Figure 1**. Residues with an average ΔrRMSF greater or less than 2 standard deviations from the mean ΔrRMSF fare considered indicators of significant changes

**Figure 1.**
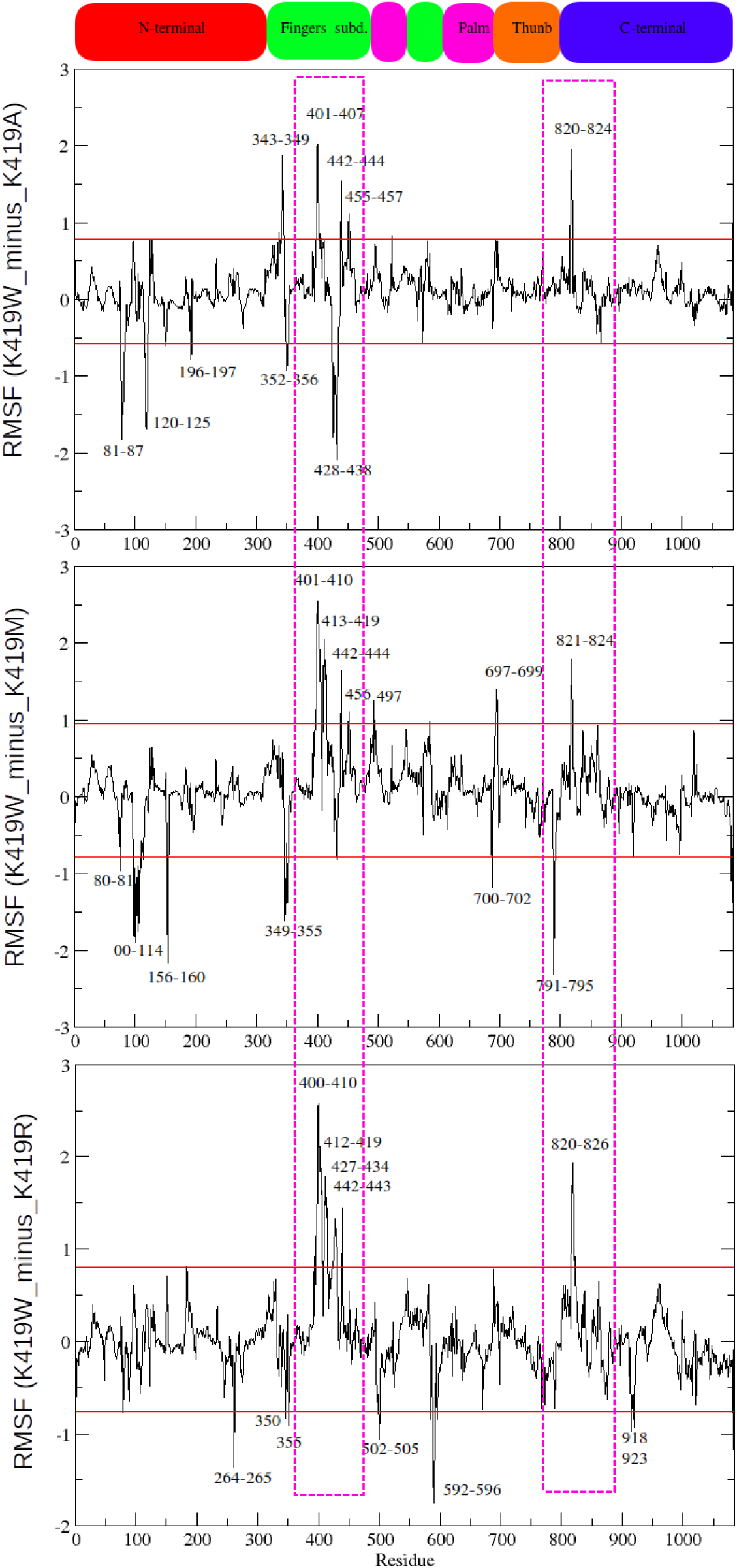
Difference in root mean square fluctuations (ΔrRMSF) of backbone atoms in K419W, compared to K419A, K419M, and K419R. The fluctuations of K419W were shown by horizontal lines (red). RNA recognition site on the VP1 protein was shown by green horizontal line.

During MD simulation run, the fluctuation regions were distributed along the protein sequence. However, the most pronounced fluctuation was observed mainly for Finger subdomain (amino acid residues 400-500, mainly 400-410) and C-terminal (mainly amino acid residues 820-824) constituted by fingers, palm, and thumb subdomains.

To statistically identify the significant collective mode of atomic motions for K419 mutants, we performed PCA on the MD trajectory. Our results showed that in K419 mutants, the overall dynamic can be described by the first three eigenvectors. To map the motion onto the structure, we therefore derived RMSF from the protein backbone by considering the first three eigenvectors. Compared to K419A, K419M, and K419R, amino acid residues mapped from K419W on RMSF data showed fluctuation at residue ranges 398-414, 441-451, and 819-823.

From the extreme structures (representing the maximum projection from the average structure) generated from eigenvector-1, eigenvector-2, and eigenvector-3, the most pronounced fluctuations were observed mainly for amino acid residues 400-406 of the K419W mutant, compared to K419A, K419M, and K419R. They shift by up to 8-15 Å inwards and outwards in the bottleneck site of the entry tunnel (**Video S2**).

Furthermore, the effect of residue fluctuations during MD simulations could be calculated from the trajectory of the changes in distance between residues located in the narrow bottleneck of the entry tunnel (**Figure 2**). Residues S401, R701, G450, and S841 were chosen for distance calculation. The S401-R701 distances are reasonable and quite stable during the 75 ns simulation time at about 18 Å then the distance decreases for both K419A and K419W mutants with the most pronounced reduction observed for K419W mutant (5 Å) until 100 ns (**Figure 2A**). The G450-S841 distances are quite stable, ranging between 18 Å and 23 Å (**Figure 2B**). The h-bond analysis of the region located between residues 398 and 414 showed different patterns and time occurrence for K419A, K419M, K419R, and K419W.

**Figure 2.**
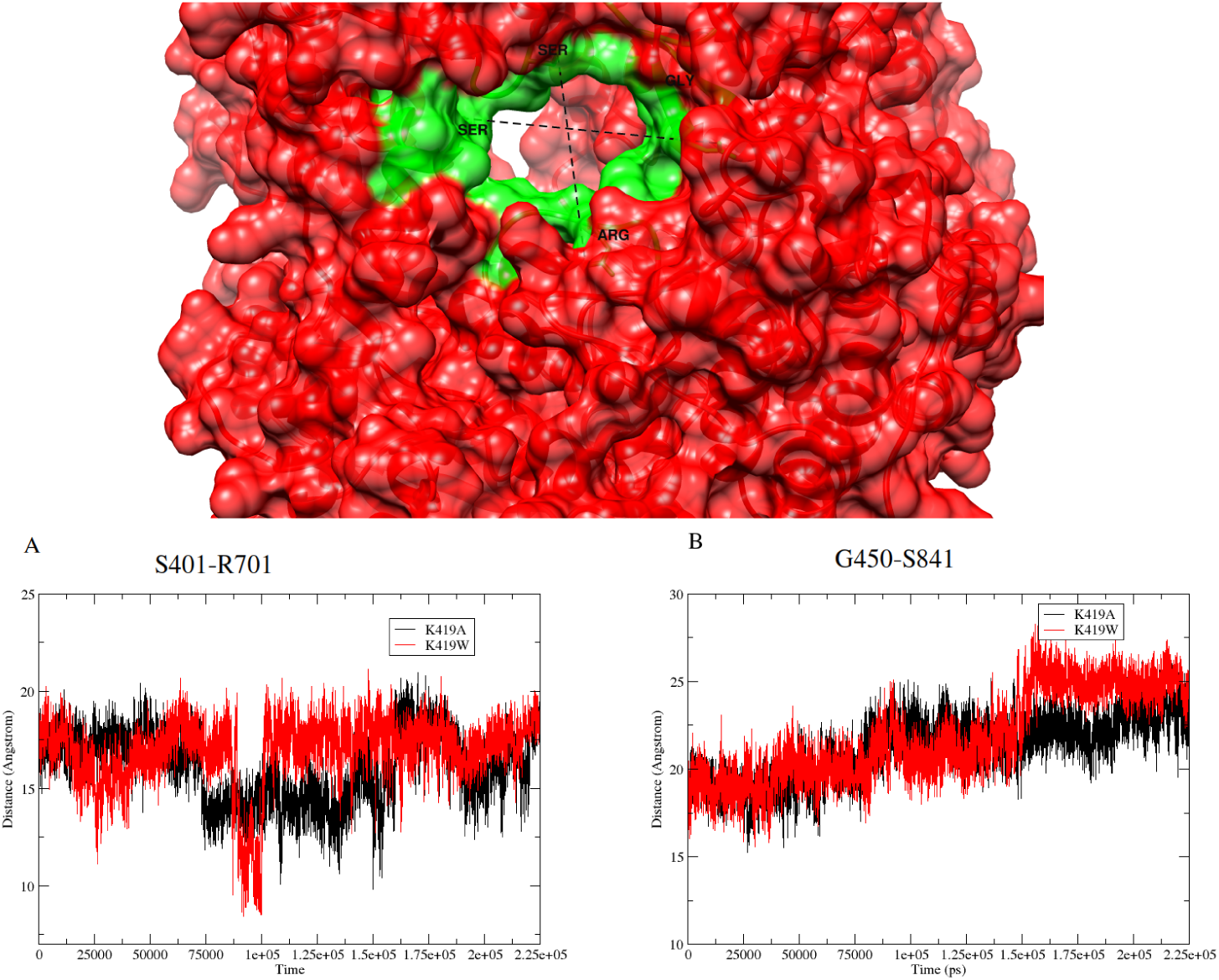
Plot of the distance between residues of the bottleneck of the RNA entry tunnel in the MD trajectories for 225 ns (green surface). (A) Distance between S401 and R701 and (B) Distance between G450 and S841.

We selected residues that may have effect on the RNA entry through the bottleneck during transcription and replication. The residue Glu404 is located in the static native structure at distances 21 Å and 25 Å from residues Arg451 and Arg452, respectively (**Figure 3**) and did not show a significant variation during MD simulations. However, the h-bond analysis of K419W mutant showed interactions between Glu404 and Arg451 with 62% occurrence and Arg452 with 54% occurrence. These h-bond interactions occur when residues move toward each other. As a result, the bottleneck of the entry tunnel becomes narrower.

**Figure 3.**
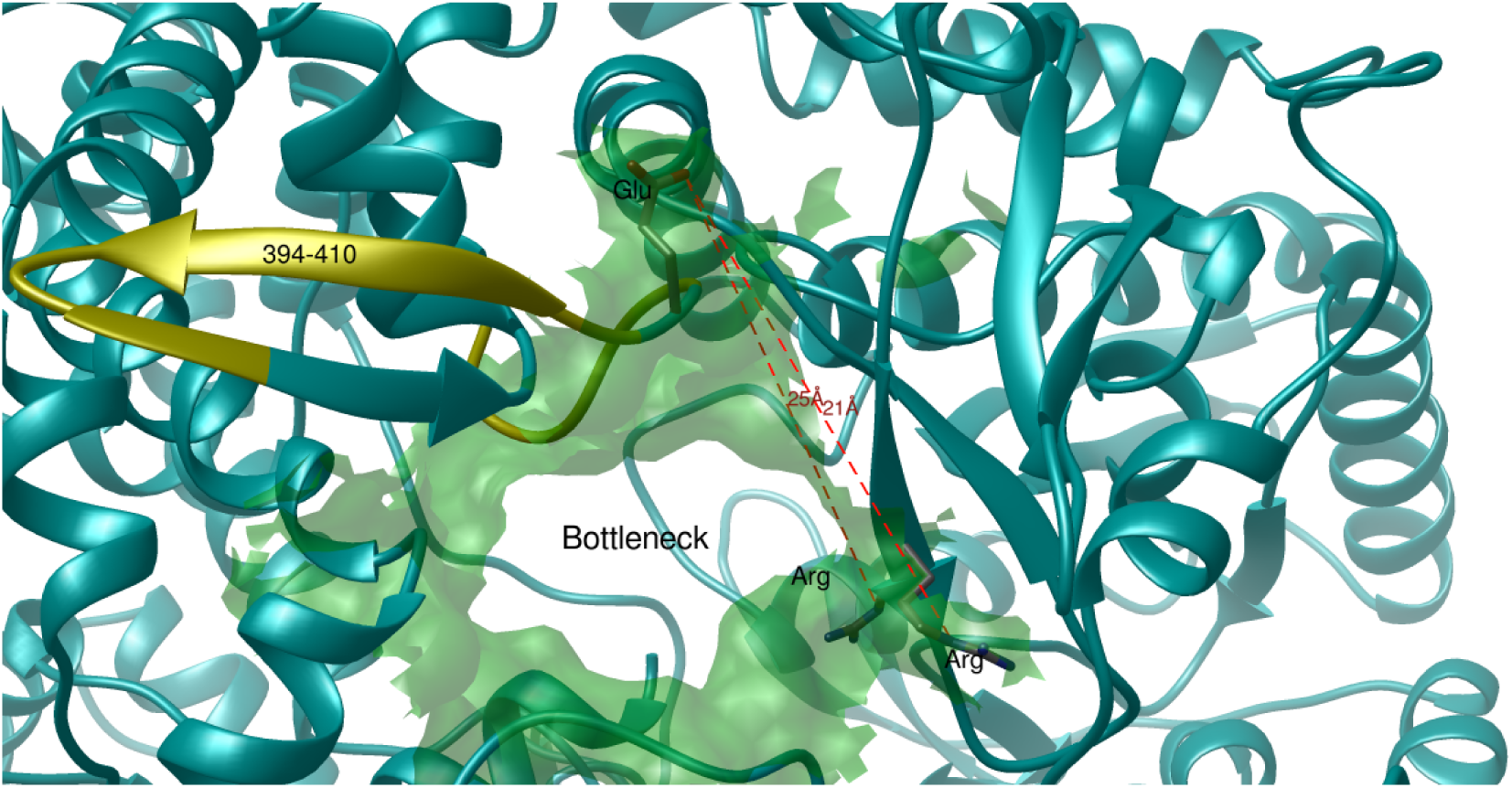
Structure representation of the RNA entry tunnel of the Native structure of VP1 protein. The bottleneck of the RNA entry was shown in green. Distances of Glu404 with Arg451 and Arg452 were shown by dash lines. The region showing the most pronounced fluctuation upon K419W mutation was shown (yellow).

Structural and functional studies have shown that VP1 protein is formed by four tunnels; one operates as the template entry tunnel, another as the nucleotide/pyrophosphate (NTP/PPi) exchange tunnel, and two others as RNA exit tunnels [8,32]. One RNA exit tunnel is used for release of ss(-)RNA template from the polymerase and is the same tunnel used for release of the dsRNA product during replication (dsRNA/ss(-)RNA exit tunnel); whereas the other RNA exit tunnel is used for release of newly made ss(+)RNAs during transcription and represents a conduit that directs nascent transcripts out of the core.

To better analyze the effect of these fluctuations on the overall topology of VP1 tunnels, we used CAVER 2.0 for the calculation of tunnels for K419A and K419W mutants. For both systems, 51 snapshots were analyzed. The initial starting point of the tunnel search was specified by atoms 4347 and 4362 of K419A and K419W, respectively. The tunnel search was performed using a maximum distance (Å) of 5 for K419W or 6 for K419A, min. probe radius (Å) of 1.5, desired radius (Å) of 5, clustering threshold (Å) of 3.5, shell depth (Å) of 4, and shell radius (Å) of 3. Redundant tunnels in each frame were removed using a linear transformation coefficient of 1 and a threshold of 1. Clustering of the tunnels identified during each simulation was performed by hierarchical average link clustering based on the pairwise distances of the tunnels computed using a linear transformation coefficient of 1 and a distance threshold of 3.5.

The results, as expected, showed that both structures present 4 tunnels: RNA entry, transcript exit, dsRNA/(-)RNA exit, and NTP entry [33] (**Figure 4**).

**Figure 4.**
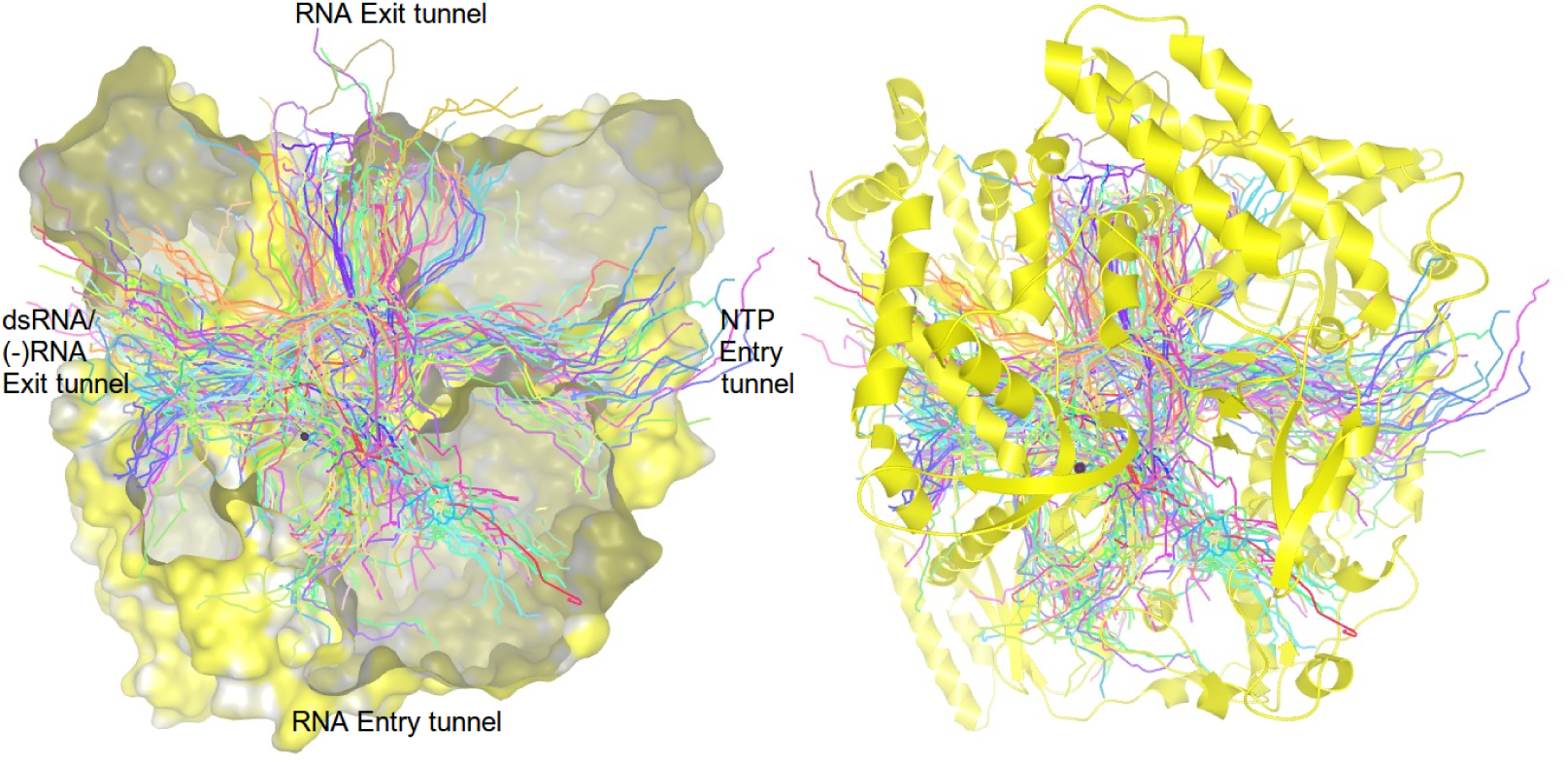
The ribbon and surface representations of VP1 protein showing the main four tunnels.

#### 3.1.2. Contribution of hydrophobic residues of motif F to dsRNA synthesis

In the molecular dynamic simulation run of three isoleucine mutations (I461, I462, and I464), the most pronounced fluctuation regions shared by these mutants were observed for residues range 516-531 and 597-603 (**Figure 5A** and **Figure 5B**). Amino acid residues 516-531 are located in the A motif (514-527); whereas amino acid residues 597-603 are located in the B motif (591-601). Our PCA results showed that fluctuations at residues 516-531 and 597-603 are more pronounced for I462A and I464A mutants than I461A, for which the most pronounced fluctuation is located around residues 520-523 and 598-600 (data not shown).

**Figure 5.**
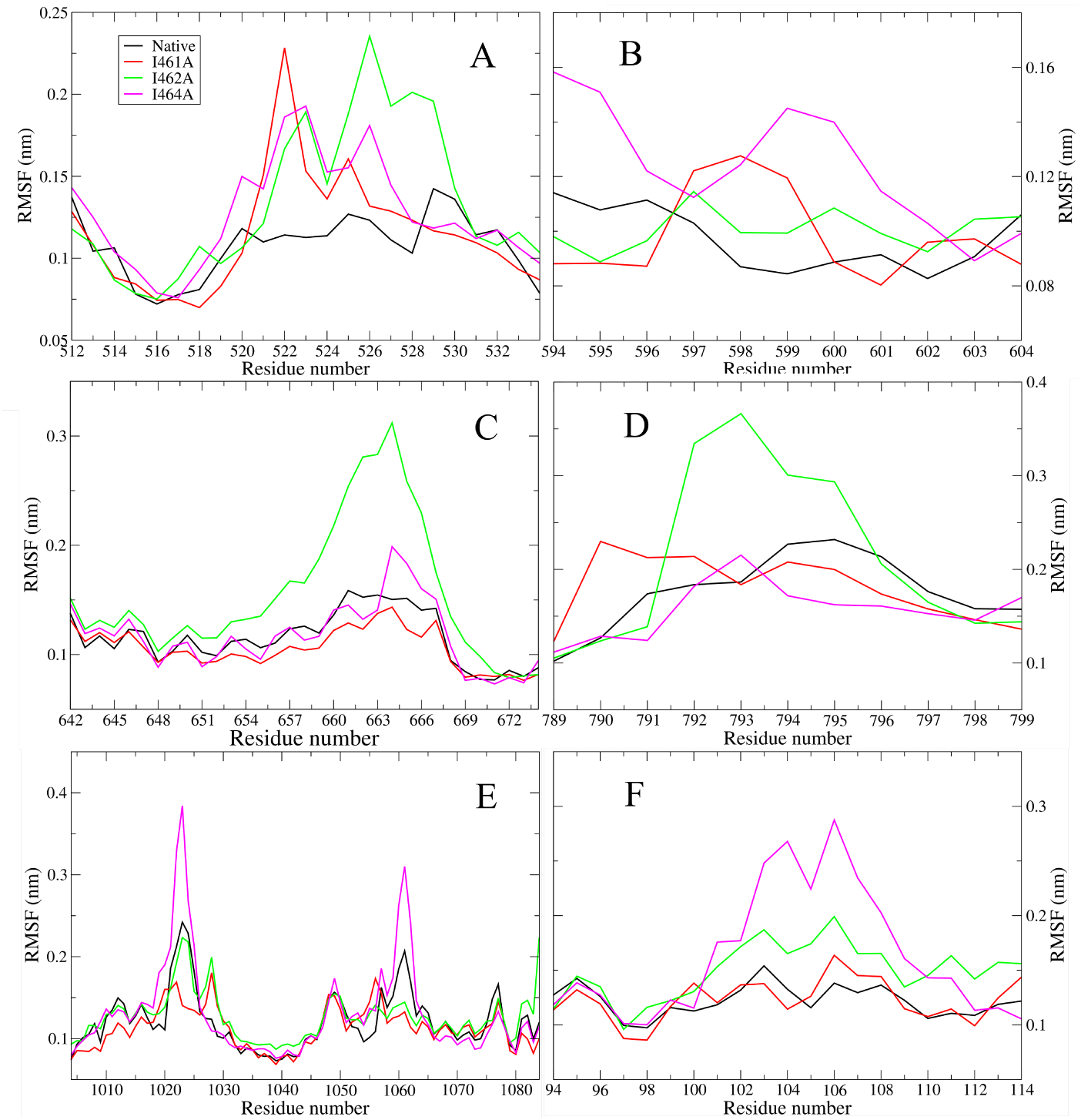
Root mean square fluctuation (RMSF) values of Cα atoms of the Native and mutants at hydrophobic residues of the motif F (I461A, I462A, and I462A) calculated during 225 ns simulation run. (**A**) and (**B**) RMSF values of fluctuations shared by the three mutants; (**C**) and (**D**) RMSF values of the additional pronounced fluctuation regions of I462A; (**E**) RMSF values of the additional pronounced fluctuation regions of I464A, (**F**) RMSF values of the additional pronounced fluctuation regions of I462A and I464A.

The mutant I462A showed additional pronounced fluctuation regions at position 644-671 (**Figure 5C**), containing the motif D (657-661), and at position 791-796 (**Figure 5D**), flexible loop that lines one side of the dsRNA/(-)RNA exit tunnel.

The mutant I464A showed additional pronounced fluctuation regions at position 1016-1026 and its juxtaposed region at position 1057-1063 (**Figure 5E**). These regions are located in the VP2 recognition site [34,35]. Moreover, additional fluctuation regions were registered for I462A and I464A mutants at positions 100-112 (**Figure 5F**).

#### 3.1.3. The effect of multiple VP1 mutation on dsRNA synthesis

Single (N186A; named N186A_1), double (N186A and F416A; named N186A_2), and triple (N186A, F416A, and R190A; named N186A_3) mutants produced different levels of dsRNA during initiation assay, from equivalent (single mutant) to significantly reduced yields (double and triple mutants), compared to native type. In addition, experimental data showed that during viral replication (initiation and elongation), double and triple mutants exhibited reductions in dsRNA synthesis of approximately 2-fold and 20-fold, respectively [10].

The MD analysis showed that the double and triple mutants shared fluctuations at residues 585-605, overlapping B motif (591-601); and residues 624-630, located in the C motif (621-639) and in the catalytic pocket near the NTP entry tunnel (**Figure 6A** and **Figure 6B**). Additional pronounced fluctuations were registered at positions 475-494 and 516-534 for N186A_2 (**Figure 6C** and **Figure 6D**) and region at positions 575-580 for N186A_3 mutant (**Figure 6E**).

**Figure 6.**
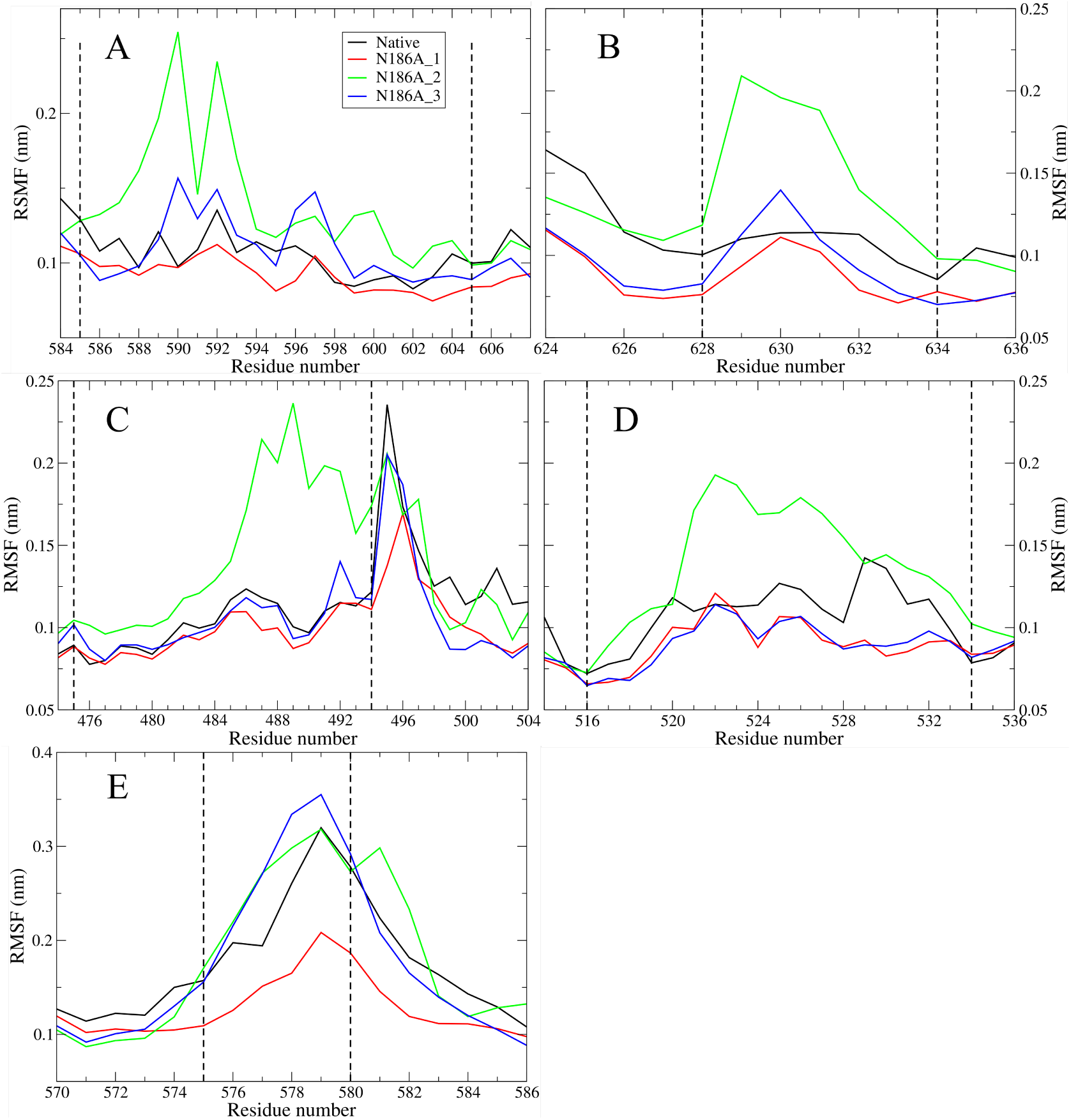
Root mean square fluctuation (RMSF) values of Cα atoms of the Native and multiple mutants (N186A_1, N186A_2, and N186A_3) calculated during 225 ns simulation run. (**A**) and (**B**) RMSF values of fluctuations shared by double and triple mutants; (**C**) and (**D**) RMSF values of the additional pronounced fluctuation regions of N186A_2; (**E**) RMSF values of the additional pronounced fluctuation regions of N186A_3.

#### 3.1.4. The contribution of finger subdomain residues of VP1 to dsRNA synthesis

The MD analysis of residues located in the finger subdomain of VP1 protein, S398A and K420A, showed fluctuations at residues 520-524 and 589-601 (**Figure 7A** and **Figure 7B**), yet supported by at least 5 eigenvectors using the PCA analysis (**Figure 7C** and **Figure 7D**), respectively.

**Figure 7.**
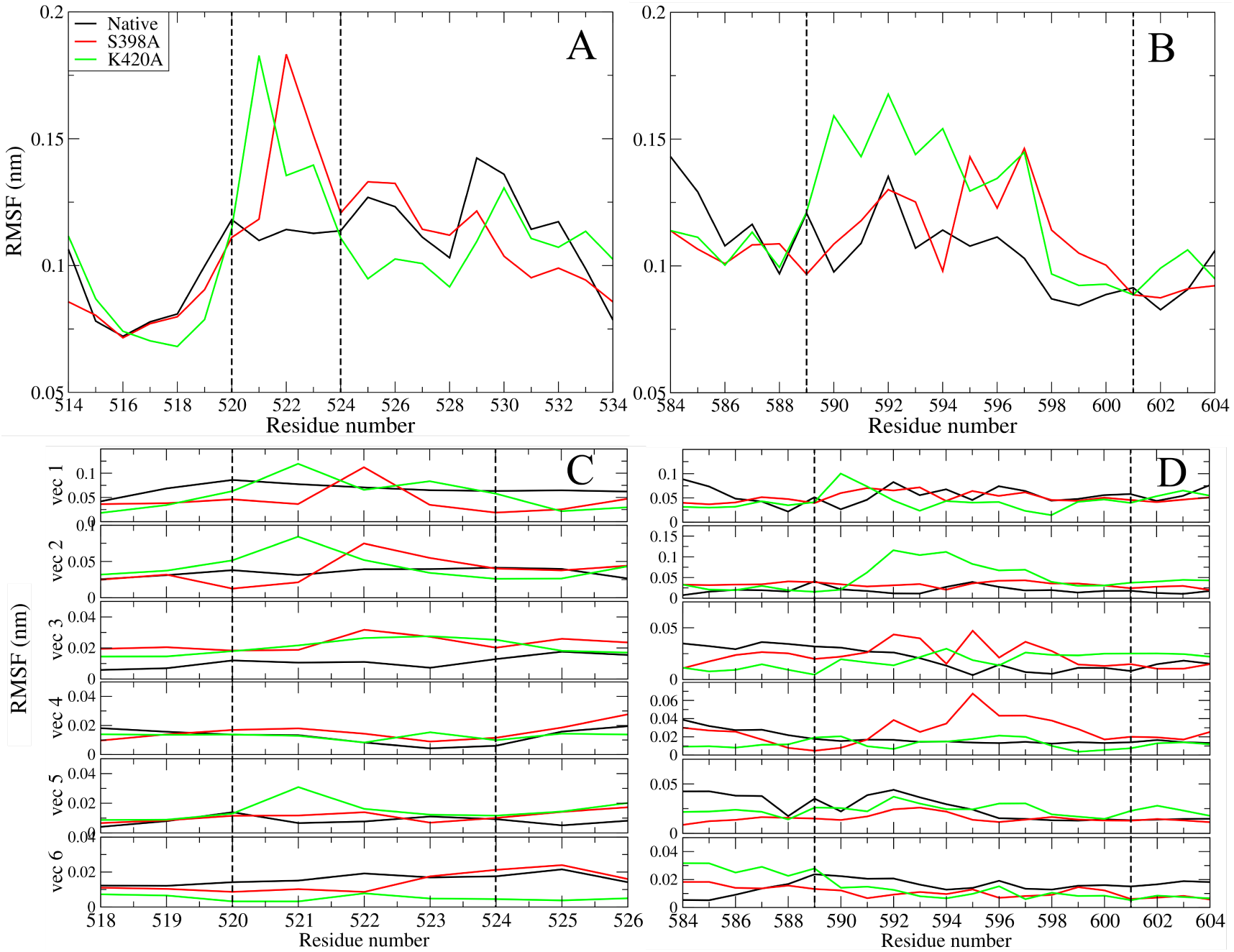
Root mean square fluctuation (RMSF) values of Cα atoms of the mutants at the finger subdomain (S398A and K420A) calculated during 225 ns simulation run (**A** and **C**). Contribution of each Cα atom to the total RMSF along the first six eigenvectors (**B** and **D**).

In addition, The PCA analysis showed evidence of pronounced fluctuation for K420A and S398A mutants in the N-terminal region (amino acid residues 1-200) (**Figure S2A**), with the most pronounced fluctuations at residues 30-60 and 97-117 for S398A and 119-127 for K420A (**Figure S2B** and **Figure S2C**). Additional fluctuations at residues range 1017-1025 for S398A mutant (**Figure S3A**) yet supported by the PCA analysis (**Figure S3B**). These residues are located in the VP2 recognition site [34] whereas residues range 97-117 and 119-127 line one side of the NTP entry tunnel. Residue 30-60 are located to the helix-loop-helix subdomain (HLH).

#### 3.1.5. Analysis of molecular dynamics simulation of (ts) mutants

It was shown that L138P mutant is localized to viroplasm and interacts with NSP2 and/or NSP5 at 31°C, but abrogates this interaction at 39°C and reduces the enzymatic activity of the polymerase *in vitro*. The percentage of dsRNA made by VP1_L138P_ at 39°C was 65% of that made at 31°C [12]. To gain insight into a possible temperature-induced structural changes resulting from the L138P mutation, MD simulations were performed. Because it was shown that the enzymatic activity reduced in a temperature-dependent manner, we choose to extend the temperature frame from 27°C to 39°C to better reveal any structural difference that may affect the VP1 activity. Structure of L138P mutant was simulated for 225 ns at 300K and 312K.

We performed a PCA on the MD trajectory to statistically identify the significant collective mode of atomic motions for L138P mutant. Our PCA result showed that the overall dynamic can be described by the first six eigenvectors. To map the motion onto the structure, we derived RMSF from the protein backbone by considering the first six eigenvectors. The results showed that amino acid residues mapped from L138P_312K_ on RMSF data showed fluctuation at residue ranges 431-433, 482-499, and around residue 746, compared to L138P_300K_ (**Figure S4**).

Additionally, the time evolution of the secondary structure of L138P at two different temperatures (300K and 312K) showed modification in the priming loop (aa 486-507) [9], mainly amino acid region 491-497 of L138P_312K_, resulting the conformation change from loop to α-helix shape between 40-105 ns of the trajectory (**Figure 8**).

**Figure 8.**
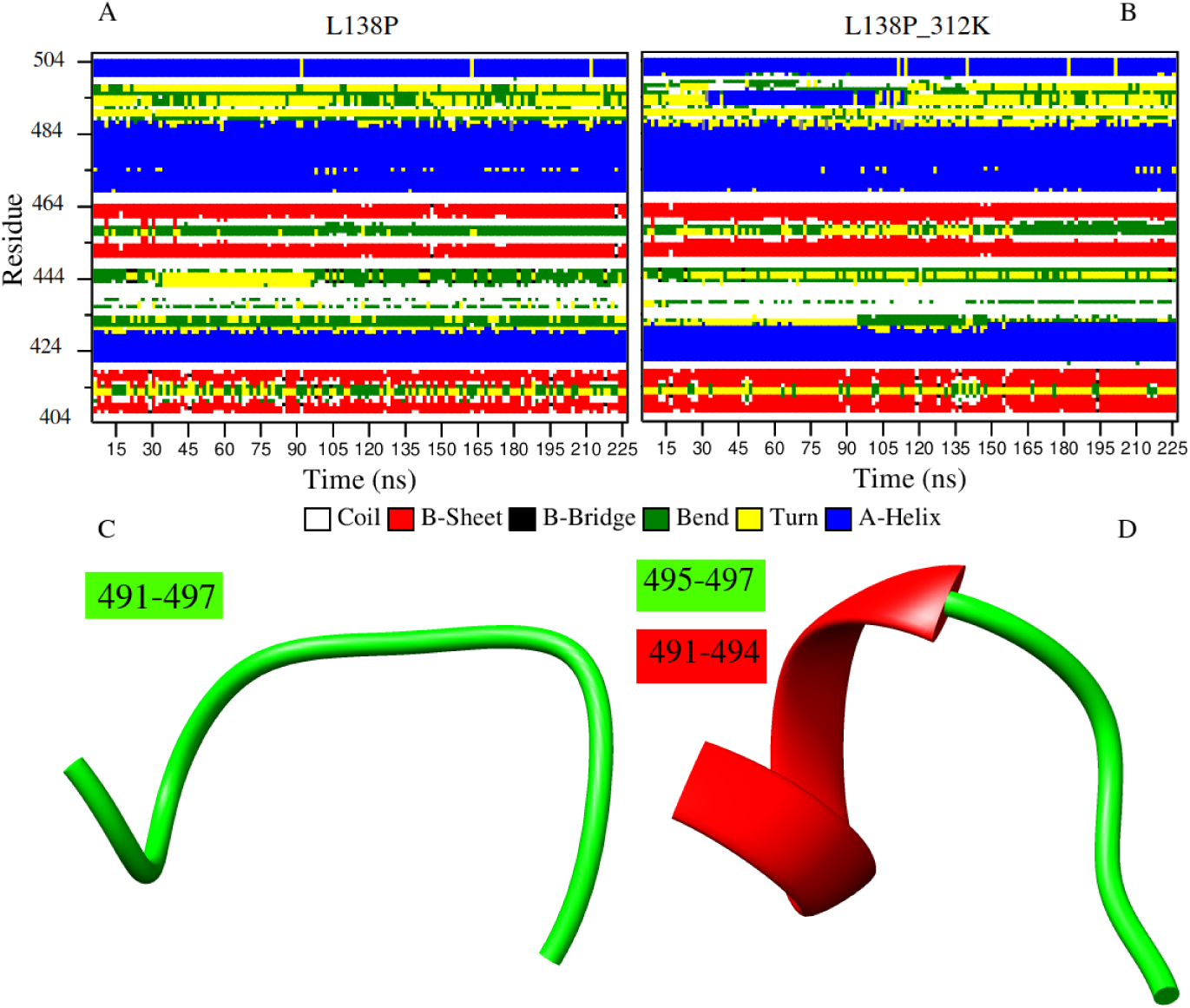
Secondary structure evolution of the L138P mutant at 300 K (**A**) and 312K (**B**). The snapshot structures of L138P_300K_ (**C**) and L138P_312K_ (**D**) extracted from Gromacs trajectories at 75 ns, in which the secondary structure is according to DSSP analysis. The regions of the loop (green) and the α-helix regions (red) are shown.

### 3.2. Average Shortest Path (L) and Betweenness Centrality (BC)

L measures the accessibility of a residue within a protein, while average L refers to the mean protein residue accessibility across all MD frames. BC and L values and their respective averages were calculated using *calc_network.py* and *avg_network.py* scripts for the Native and mutant proteins. Thereafter, average ΔrL (mutant minus Native) values were calculated for each Native and mutant protein system across all MD frames (**Figure S5**). A decrease to ΔrL indicates that residues in mutants are moving closer to each other with respect to the Native and becoming more accessible, whereas, an increase to ΔrL indicates a decrease in residue accessibility within the mutant in comparison to the Native. Residues with an average ΔrL greater or less than 2 standard deviations from the mean ΔrL for each protein system are considered indicators of significant changes to accessibility. The results showed that ΔrL calculations identified subtle differences.

For highly dynamic proteins, it was reported strong positive correlation between L and RMSF [36,37], and moderate and weak inverse correlation between BC and L or RMSF, respectively [37]. These findings were partially supported in the present study by calculating the pairwise Pearson’s correlation coefficient of average L, average BC, and RMSF for each VP1 structure (**Table 2**). The average L and RMSF were weakly to moderately correlated (r range=0.29– 0.59); the average BC and RMSF^−1^ were moderately correlated (r range=0.25–0.42); the average BC and L^−1^ were highly correlated (r range=0.77–0.81); whereas BC and L or RMSF showed high (r range between –0.74 and –0.77) and weak to moderate (r range between –0.19 and –0.38) negative correlation, respectively.

**Table 2.** Pearson’s correlation coefficient values: Pairwise comparisons of average BC, average L and RMSF.

|  | L vs RMSF (225ns) | BC vs RMSF (225ns) | L vs BC | L<sup>-1</sup> vs BC | RMSF<sup>-1</sup> (225ns) vs BC |
| --- | --- | --- | --- | --- | --- |
| I461A | 0.44 | -0.34 | -0.75 | 0.78 | 0.41 |
| I462A | 0.36 | -0.26 | -0.77 | 0.81 | 0.33 |
| I464A | 0.41 | -0.28 | -0.76 | 0.79 | 0.35 |
| K419A | 0.48 | -0.33 | -0.74 | 0.77 | 0.40 |
| K419M | 0.42 | -0.34 | -0.76 | 0.79 | 0.40 |
| K419R | 0.29 | -0.19 | -0.76 | 0.80 | 0.25 |
| K419W | 0.32 | -0.22 | -0.74 | 0.78 | 0.26 |
| L138P | 0.43 | -0.27 | -0.77 | 0.80 | 0.31 |
| L138P<sub>312K</sub> | 0.44 | -0.31 | -0.76 | 0.80 | 0.38 |
| S398A | 0.47 | -0.35 | -0.76 | 0.79 | 0.40 |
| K420A | 0.59 | -0.38 | -0.75 | 0.79 | 0.42 |
| N186A_1 | 0.47 | -0.31 | -0.75 | 0.78 | 0.35 |
| N186A_2 | 0.50 | -0.35 | -0.75 | 0.78 | 0.39 |
| N186A_3 | 0.40 | -0.24 | -0.77 | 0.80 | 0.28 |

The results indicate that the majority of changes to residue accessibility for the most of mutants, with exceptions, occur mainly in the N-terminal, Finger, and the C-terminal domains (**Figure S5**). For K419 mutants (K419A, K419M, K419R, and K419W), we noted that the global distribution of average ΔrL values across mutants was generally similar. Although all mutants showed similar profiles, K419W shows increase to residue accessibility between residues 404 and 407 and decreases to residue accessibility between residues 584 and 586; these regions are surface exposed and close to the bottleneck tunnel of the VP1 protein (**Figure S5A**).

The I461A, I462A, and I464A mutants showed different average ΔrL patterns. I461A mutant shows more pronounced decrease to residue accessibility in the N-terminal domain, when compared to I462A and I464A mutants for which an additional decrease to residue accessibility was shown for residue 107; yet more pronounced for I464A mutant (**Figure S5B**). The results showed increase to residue accessibility for I461A and I462A between residues 185-190, but not for I464A mutant. The decrease to residue accessibility in the Finger subdomains was more pronounced for I464A mutant, mainly for residues 441-446 and 464-467. Additional decrease to residue accessibility in Palm subdomain was shown for I461A (aa 522) and I462A (aa 662) mutants. I461A mutant shows more pronounced decrease to residue accessibility in the C-terminal domain, when compared to I462A and I464A mutants. Additional decrease to residue accessibility was shown for residues 790-796 of I462A mutant.

The analysis of single (N186A_1), double (N186A_2), and triple (N186A_3) mutants showed different average ΔrL profiles (**Figure S5C**). The decrease to residue accessibility in the N-terminal domain was more pronounced for N186A_2 mutant, mainly for residues 111 and 121-126; whereas N186A_3 mutant shows more pronounced increase to residue accessibility for residues 127-133 and 185-191 of the N-terminal domain. The decrease to residue accessibility in the Finger subdomains showed different pattern for the three mutants. While N186A_1 and N186A_2 mutants show similar profile for the the first Finger subdomain (residues 333 to 488), N186A_2 and N186A_3 mutants show more pronounced decrease to residue accessibility between residues 577 and 581 of the second Finger subdomain (residues 524 to 595). Minor changes of residue accessibility were shown for N186A_2 and N186A_3 mutants in the C-terminal domain, when compared to N186A_1 mutant.

The K420A and S398A mutants showed slightly difference in N-terminal domain, mainly a decrease in residue accessibility for residues 121-128 (K420A mutant) and residue 238 (S398A mutant). Although similar changes were shown for both mutants, K420A mutant shows decrease to residue accessibility for residues 1020-1023 in the C-terminal domain (**Figure S5D**).

Finally, the analysis indicates that all changes to residue accessibility for L138P mutant are distributed in all protein domains (**Figure S5E**), except the Thumb domain. The decrease to residue accessibility for L138P mutant was shown mainly for residues 238, 347, 351, 433, 577-586, and 1023 which was not observed for L138P_312K_ mutant for which the most pronounced decrease to residue accessibility was shown for residues 789-793 in the C-terminal domain.

During MD simulation, protein residues are constantly communicating with one another, and BC allows for identification of the most important residues for protein structure and function. Residues with the highest BC are most important for protein communication. When mutations are present, residue communication could change, and analysis of these changes would allow for the identification of compensatory mechanisms employed by mutants in an attempt to maintain protein function and stability.

To observe the BC differences between the Native and mutant proteins, average ΔrBC (mutant minus Native) was calculated (**Figure S6**). A decrease to ΔrBC indicates a decrease in residue usage within the mutant whereas an increase to ΔrBC demonstrates increased residue usage. As for ΔrL, residues with an average ΔrBC greater or less than 2 standard deviations from the mean ΔrBC for each protein system indicating significant changes to residue communication.

The results indicate that the majority of changes in residue usage for the most of mutants occur mainly in the central polymerase domain (Fingers, Palm, and Thumb subdomains) and the C-terminal domain.

K419W shows a high ΔrBC value for residues 419 of the Finger subdomain, Compared with the remaining K419 mutants, suggesting that this residue plays an important role for communication in K419W mutant. In addition, the low ΔrBC value for the residue 453 of the same mutant shows a decrease in residue usage, also when compared to the remaining K419 mutants. Furthermore, residues in the C-terminal domain of the K419W mutant were shown to be less important (residues 922 and 991-999) as they showed low BC values, also compared with K419A, K419M, and K419R mutants (**Figure S6A**).

Although the I461A, I462A, and I464A mutant showed similar ΔrBC profiles, few residues in the Finger subdomains showed different ΔrBC values (**Figure S6B**), mainly residue 338, and 591 of the I462A and I464A mutants which showed more pronounced decrease in residue usage, compared with I461A mutant. Furthermore, the I464A mutant showed increase in the residue usage at position 465.

The N186A_1, N186A_2, and N186A_3 mutants showed similar ΔrBC profiles. However, N186A_3 mutant showed some difference in residue usage during MD simulation; residues at position 128-135 and 1002 showed pronounced increase in residue usage, compared with the remaining mutants. More pronounced decrease in residue usage was shown mainly for residues 453 and 591 of the Finger subdomain. Residues 596 of N186A_2 mutant showed decrease in residue usage while the same residue showed increase in residue usage for N186A_1 and N186A_3 mutants (**Figure S6C**).

Although the K420A and S398A mutants showed similar residue usage, the residue 498 of the Palm subdomain was shown to be more important in the K420A mutant. Pronounced change in residue usage was shown for residues 591 and 594 (Finger subdomain) of S398A mutant, showing decrease and increase in residue usage, respectively (**Figure S6D**).

The main difference in ΔrBC profiles of L138P and L138P_312K_ was shown for residue 249 of L138P mutant, showing increase in residue usage and an increase in residue usage for residue 701 (Thumb subdomain) of L138P_312K_ mutant (**Figure S6E**).

### 3.3. Residue contact map

Contact map analysis was performed to identify mutation associated short-range changes to the protein network (**Figure S7**). Contact maps show all weighted/frequency of interactions occurring between a network of residues within 7 Å over the MD simulation. These weighted contacts include all interactions such as, but not limited to, van der Waals, hydrogen bonds and/or electrostatic interactions [27].

Analysis of K419 mutants and Native contact maps indicates no changes to interactions occurring between Lys419 and Arg419 in the Native and K419R mutant, respectively; however Ala419 in the K419A mutant shows a decrease to interactions with Ser401, whereas Met419 in the K419M mutant shows increase to interaction with Ser841, comparing to Native structure. The most variation in the contact maps was shown for K419W; where Trp419 shows decrease to interactions with Ser399 and His423. No change was shown for Arg419 of K419R mutant (**Figure S7A**).

The residue network analysis of Ileu461 of I461A mutant showed decrease to interactions with Ser591; Ile462 of I462A mutant showed increase to interactions with Ala590, Ile464, and Arg451; Ile464 of I464A mutant showed increase to interactions with Gly403 and Asn402, whereas it shows decrease to interactions with Leu449 and Gly450 (**Figure S7B**).

The results obtained for N186A_1 showed increase to interactions of Ala186 with Tyr191, whereas it showed decrease to interactions with Asp194 (**Figure S7C**). Ala416 of N186A_2 did not show difference in residue network, comparing to Phe416 of the Native structure. However, Ala186 showed a decrease to interactions with Asp194 and Lys188 (**Figure S7D**). The most pronounced changes in residue network interactions were shown for N186A_3 mutant. Upon N186A mutation, Ala186 showed an increase to interactions with Tyr191 and a decrease to interactions with Asp194 and Ala190. The latter residue, as a result of R190A mutation, showed an increase to interactions with Met130 and a decrease to interactions with His185 and Glu182 (**Figure S7E**).

Ala398 of S398A mutant showed a slightly increase to interaction with Glu1088, whereas Ala420 of K420A mutant showed an interaction with Gly403, which is absent in the Native structure (**Figure S7F**).

The Ile177 constitutes the main difference in L138P and L138P_312K_ mutants. This residue showed an increase interaction with Pro138, comparing to the Native and L138P_312K_ structures (**Figure S7G)**.

Dynamic cross correlation captures the collective behavior of the atoms, i.e., the degree to which they move together. The color-coded maps provide a visualization of the correlated motions of the structures, where warm colors (from yellow to red) indicate relatively higher positive correlation, whereas the cold color (from cyan to blue) represents relatively highly anti-correlation.

For Native and mutants, both highly correlated and highly anti-correlated residues were observed. These structures showed similar dynamic cross correlation profile (**Figure S8**).

## 4. Discussion

To explore the effect of mutations on VP1 protein, the MD simulation of the mutated VP1 at RNA entry bottleneck (K419A, K419M, K419R, and K419W) showed interesting results. The K419W mutant showed fluctuation at amino acid residues 398-414, compared to K419A, K419M, and K419R mutants. This fluctuated region is located in the RNA recognition site of VP1 protein. In addition, K419W mutant showed fluctuations at amino acid residues 441-451, located in the NSP2 recognition site [8] and close to conserved arginine residues of VP1 (Arg451, Arg452, Arg457, Arg458, and Arg460), shown for their important contribution to dsRNA synthesis [11]; and amino acid residues 819-823, located close to the NSP2 recognition site [8], which lines the side of the dsRNA product exit tunnel of VP1. Binding of NSP2 at these sites was suggested to bring it into close proximity with the 5′ (-)RNAs, leading to more efficient γ-phosphate hydrolysis [38]. The importance of the pronounced fluctuation at the amino acid residues 398-414 of the K419W mutant is supported by a recent study on the structural differences between non-transcribing and transcribing DLPs, showing that loop 397-404 of the fingers domain retracts from the central cavity to allow the (-)RNA to move into the catalytic site through a bottleneck towards the palm in transcribing VP1 [9]. Furthermore, it was reported that after the split of the dsRNA molecule during transcription, the 3′CS- (AAAAGCC-3′) of the (-)RNA strand weakly interacts with the β-sheet subdomain (residues 400-419) in the fingers [7]. The increase to residue accessibility, as reported by calculating the average ΔrL near the bottleneck tunnel of the VP1 protein is likely due to the displacement of the bottleneck residues. Residues 584-586 are located close to the NSP2-binding sites on VP1 [8] and their decrease in accessibility may affect this binding. The decrease in residue usage of residues 453, 922 and 991-999 of K419W mutant, as shown by the average of ΔrBC analysis, may affect the function of VP1 as residue 453 is located to NSP2-binding sites on VP1 (aa 442 to 458) [8] whereas residues 922 and 991-999 are located in the C-terminal domain, responsible for splitting the dsRNA product during transcription [7]. The decrease of interactions of Trp419 (K419W mutant) with Ser399 and His423, as shown by contact map analysis, is the result of the bottleneck displacement. The decrease of Ala419 interaction with Ser401 of K419A is likely to decrease the steric hindrance and electrostatic interactions with the 3’-end of the template during initiation, whereas the increase of Met419 interaction with Ser841 of K419M allows the bottleneck structure of VP1 to be wider for easily move the template into the catalytic pocket [10]. Altogether suggest that the fluctuation of residues 398-414 likely diminished VP1 replication activity by severely restricting RNA entry through the bottleneck and moderated both initiation and the kinetics of elongation during dsRNA synthesis. However, Caver analysis showed that the K419 mutants did not affect the overall topology of the protein tunnels. Moreover, the RNA entry constituted by two pathways outside the catalytic site within the RNA entry tunnel. The pathways likely constitute the two binding sites for (-)RNA and (+)RNA, as a results of an open genomic duplex in the VP1 in non-transcribing state, as reported previously [7].

VP1 polymerase contains canonical motifs A through F [33,39]. Residues in these motifs interact with templates, incoming NTPs, and the divalent metal ions (Mg2+) that catalyze phosphodiester bond formation during viral transcription and replication. Three isoleucine residues (I461, I462, and I462) located at the F motif of RdRp polymerase, except I461, are involved in sequence-independent contacts with the UGUG bases of the 3’CS+ primarily with the phosphate linkages and ribose groups of the RNA backbone (**Table 1**). Substitution of these amino acid residues by alanine showed decrease in dsRNA synthesis by 2-fold for I461A and 20-fold for I462 and I464 during replication assays [10]. Experimental data showed that for I461A mutant, dsRNA synthesis was slightly reduced during replication assays yet equivalent during initiation assays, compared to the Native VP1 protein. In contrast, I462A and I464A mutants exhibited greater reduction in dsRNA product in replication assays and highly impaired dsRNA synthesis in initiation assays [10]. The RMSF analysis showed fluctuations in the B and D motifs of these mutants. Motif B contains a number of conserved residues, with several anchoring the 3′ terminus of the (+)RNA template inside the enzyme. It was suggested a particularly important role of motif B in aligning the template strand with NTPs in the catalytic center [40]. Based on the RdRp of poliovirus, it was suggested that motif D can coordinate the export of the PPi group from the active site once catalysis has taken place, thereby triggering the translocation of the RNA [41]. In addition, NMR studies have indicated the inability of motif D to achieve its optimal conformation for catalysis when an incorrect nucleotide is incorporated, thereby demonstrating its role in the selection of NTPs [42]. Moreover, the Average Shortest Path analysis showed decrease to residue accessibility in motif F of I464A, motif A of I461A, and motif D of I462A mutants whereas Betweenness Centrality analysis showed decrease in residue usage in the N-terminal subdomain (residue 338) and the motif B (residue 591), mainly for I462A and I464A mutants. Motif A houses the catalytic motif DX_4_D in which the second aspartate together with a conserved asparagine from motif B plays a crucial role in the discrimination of NTPs over deoxy NTPs by forming a h-bond with 2’OH of the incoming NTP [43]. Additionally, the residue network analysis showed decrease of interactions between motif F residue (Ileu461) and motif B residue (Ser591) of I461A mutant and decrease of interactions within motif F residues (Ile464 with Leu449 and Gly450) of I464A mutant; the increase of interactions within motif F (Ile462 with Ile464 and Arg451), between motifs F and B (between Ile462 and Ala590) of I462A mutant, and between motif F residue (Ile464) and the bottleneck residues (Gly403 and Asn402) of I464A mutant. The residue network changes of motif F residues may be resulted from the fluctuations of B and D motifs. Motif F is formed by positively charged residues that shield the negative charges of the phosphate group of incoming NTP [43,44] and residue network stability is crucial for its function. These results suggest that fluctuations and residue network changes in the discussed above VP1 motifs may hamper their role in selecting the suitable NTP during elongation.

The RMSF analysis of the multiple VP1 mutations (N186A_1, N186A_2, and N186A_3) showed fluctuations in the B and C motifs. The later, one of the most conserved motifs, essential for binding the Mg2+ [45-48]. The additional fluctuation regions for N186A_2 mutant, residues 475-494 overlaps putative priming loop (aa 486-507) [9] and close to motif F (residues 445-466). Residues 516-534 are located in the A motif (residues 514-527), important for the coordination of Mg2+ and the interaction with the incoming NTP during viral replication, as discussed above. For N186A_3, residues 575-580 are located within NSP2 recognition site (aa 562-578) [8]. The Average Shortest Path analysis showed decrease to residue accessibility at the N-terminal subdomain of N186A_2 mutant and at the junction between A and B motifs of the Finger subdomains of N186A_2 and N186A_3 mutants, close to NSP2-binding sites on VP1 [8]. Betweenness Centrality analysis showed decrease in residue usage at NSP2-binding site on VP1 and in the motif B of N186A_3 mutant. The residue network analysis showed decrease of interactions at the N-terminal subdomain of N186A_2 and N186A_3 mutants, close to the dsRNA entry channel. As reported previously, once the VP1 is transcriptionnally active, the (-)RNA strand traverses the template entrance through a bottleneck towards the active site, and the VP1 initiates (+)RNA synthesis templated from the 3′CS-of the (-)RNA molecule [8]. Then, the (-)RNA pairs with complementary NTP in the active site between the fingers and the palm. The incoming NTPs are in position to form a backbone with the 5′ end of the nascent (+)RNA [7]. It was shown that the conserved acidic residues of motif A (Asp520 and Asp525) and C (Asp631 and Asp632) are involved in coordinating two Mg2+ ion in the catalytic site [40,49,50]. In addition, it was reported that two additional residues of motif A (S522 and Q523) are positioned suitably for interaction with the γ-phosphate of an incoming NTP. In addition, according to recently reported structure of transcribing VP1 [PDB ID: 6OGZ], residues Thr596, Asn600, Arg452, Arg457, Arg591, Arg460, Asp520, Asp525, Trp524, Gln523, Asp631, and Asp632 interact with the incoming NTP and the nascent (+)RNA molecule during elongation [7]. Indeed, significant fluctuations and loss of interactions among residues at VP1 motifs may affect the transcription initiation and elongation of template.

Mutations of residues at the finger subdomain showed decrease in dsRNA synthesis by 2-fold and 20-fold for S398A and K420A mutants during replication assays, respectively [10]. Furthermore, experiments showed that S398A mutant highly reduced dsRNA synthesis during replication assays yet equivalent to Native VP1 protein during initiation assays, compared to Native VP1 protein. In contrast, K420A mutant exhibited greater reduction in dsRNA product in replication assays and highly impaired dsRNA synthesis in initiation assays [10]. Residues S398 and K420, among others (Ser401, Lys419, Gly592, Lys594, and Lys597), are shown to be involved in anchoring the 3′CS+(5′-UGUGACC-3′) of (+)RNA into the template entry tunnel during replication/packaging [8]. These finger residues interact, via H-bonds, with the phosphate linkages (Ser398 and Lys419) or riboses (Lys420) of the RNA backbone [10]. These mutations showed RMSF fluctuations in the A and B motifs, important for viral replication, as discussed above. S398A mutant showed significant fluctuations at the N-terminal of VP1 protein. Recently, it was reported that during the initiation of the transcription step (dsRNA recognition), the HLH subdomain of the N-terminal (residues 31–69, HLH subdomain) splits the dsRNA encased within the core by binding to the 5′-cap-end (m7G(5′)ppp(5′)GGC) of the (+)RNA [7]. The Average Shortest Path analysis showed decrease to residue accessibility in the N-terminal (K420A and S398A mutants) and in the C-terminal subdomains (K420A mutant). Betweenness Centrality analysis showed decrease in residue usage in the B motif of S398A mutant. The residue network analysis showed a new interaction between Ala420 and a residue of the bottleneck (Gly403), may be as a result of the fluctuations of A and B motifs.

Finally, the MD simulation applied for L138P mutant at two different temperatures showed RMSF fluctuations mainly at residues of motif F and priming loop regions of L138P_312K_. The latter is suggested to stabilize the P-site NTP in the catalytic pocket during initiation [11]. Polymerization is a highly dynamic process, and loops are commonly flexible elements within protein structures. The conformation change of this flexible loop, as shown for L138P_312K_, to a less flexible α-helix shape may affect the polymerization process, mainly to maintain the P-site NTP that needs stabilization in the catalytic pocket during initiation. The Average Shortest Path analysis showed pronounced decrease to residue accessibility at the C-terminal of L138P_312K_. Betweenness Centrality analysis showed increase in residue usage at NTP/PPi entry channel (residue 249) of L138P, compared to L138P_312K_ mutant for which residue residue at the bottleneck tunnel (residue 701) was shown important.

During early viral replication, viroplasms constitute the specific structures of the cell cytosol for the assembly of progeny cores, formed mainly by NSP2 and NSP5 [52,53] with poorly defined interaction interfaces with VP1 [51,38,54,55]. We could not suggest a relationship of the obtained results with the experimental data about the viroplasm formation using L138P_312K_ because no experimental data about the NSP5 recognition sites on VP1 are available.

## 5. Conclusions

We carried out MD simulations to analyze the effects of VP1 mutations in the template entry tunnel and catalytic site on the viral dsRNA synthesis which may provide insight into initiation and viral replication processes. Although the MD simulations fluctuations in different parts of the protein structure, the most pronounced fluctuations were registered mainly in residues located in the finger subdomain, between the two palm subdomains (aa 515-535 and aa 585-605) (**Figure S9**). These regions, the most sensitive to VP1 mutations, are located in the catalytic site responsible for RNA recognition and NTP interaction during RNA transcription and replication. The exception was for K419W which showed the most pronounced fluctuations in the RNA entry bottleneck. The present work helps to get better insight in rotavirus VP1 function by analyzing the structural behaviors of mutations in the RNA entry tunnel of rotavirus VP1 using MD simulations and it paves the route for the rational design of antivirals, as they might complement the available vaccines, frequently challenged by the prevalence of new reassortment strains.

## Supporting information

Supplementary materials of the article.

## Author Contributions

Conceptualization, N.A.; methodology, N.A., M.S. and G.C.; software, N.A. and G.C.; resources, N.A. and G.C.; writing—original draft preparation, N.A. and G.C.; writing—review and editing, N.A., M.S. and G.C. All authors have read and agreed to the published version of the manuscript.

## Acknowledgments

This work was supported by the Tunisian Ministry of Higher Education and Scientific Research and by Fulbright fellowship Program (number G-1-00005) at the University of Florida, USA, and by the ‘Departments of Excellence-2018’ Program (Dipartimenti di Eccellenza) of the Italian Ministry of Education, University and Research, DIBAF-Department of University of Tuscia, Project ‘Landscape 4.0–food, wellbeing and environment’. Marco Salemi is supported, in part, by the Stephany W. Holloway University Chair. We acknowledge Cineca and ELIXIR-IIB for computing resources.

## Conflicts of Interest

The authors declare no conflict of interest.

