## Supplementary figures and images for "New Insights Into the Effect of Residue Mutation on the Rotavirus VP1 Function Using Molecular Dynamic Simulations"

### Figure S1.png

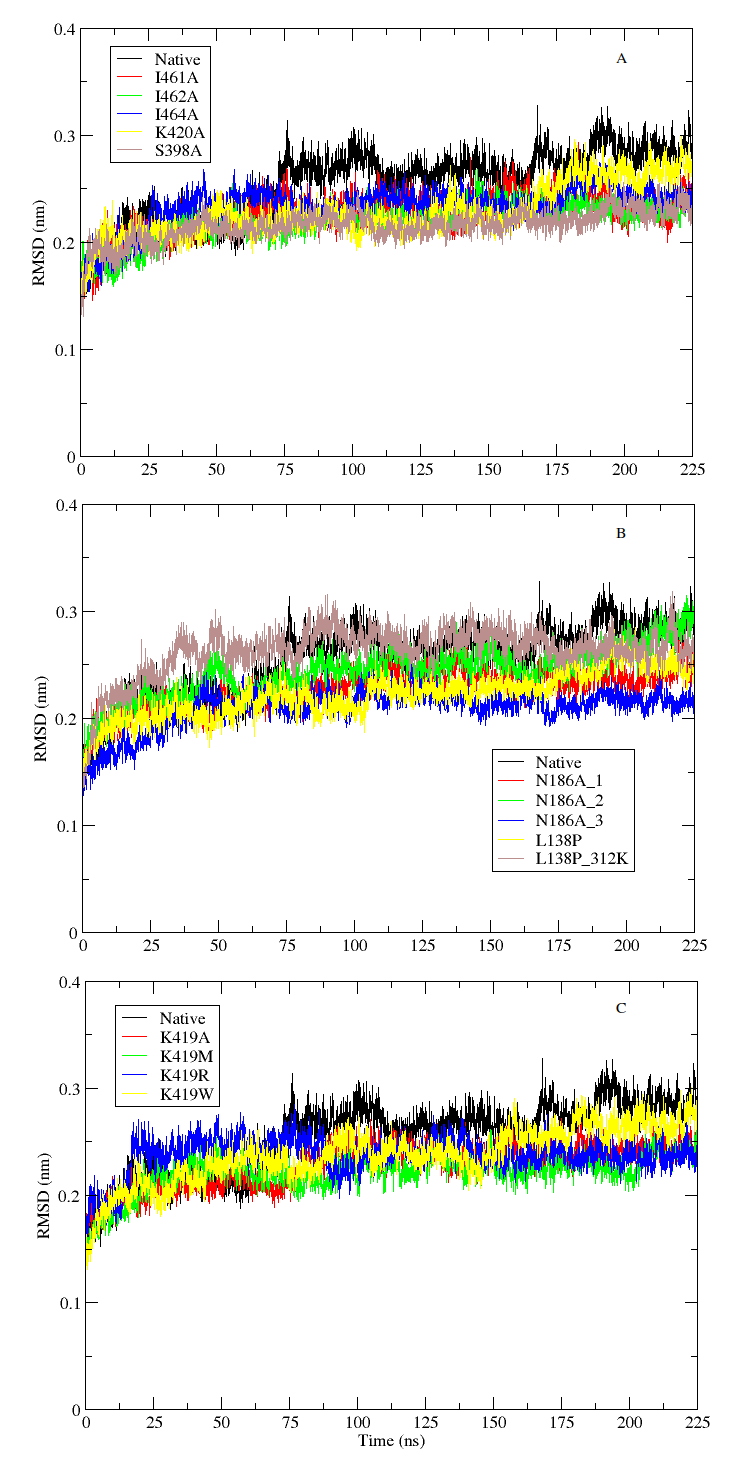

### Figure S2.png

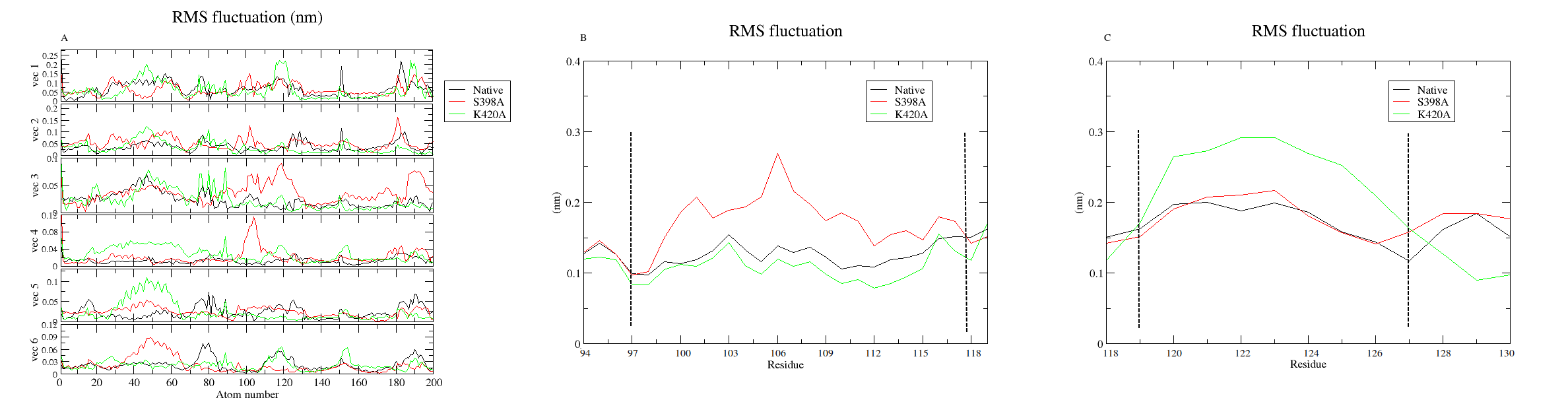

### Figure S3.png

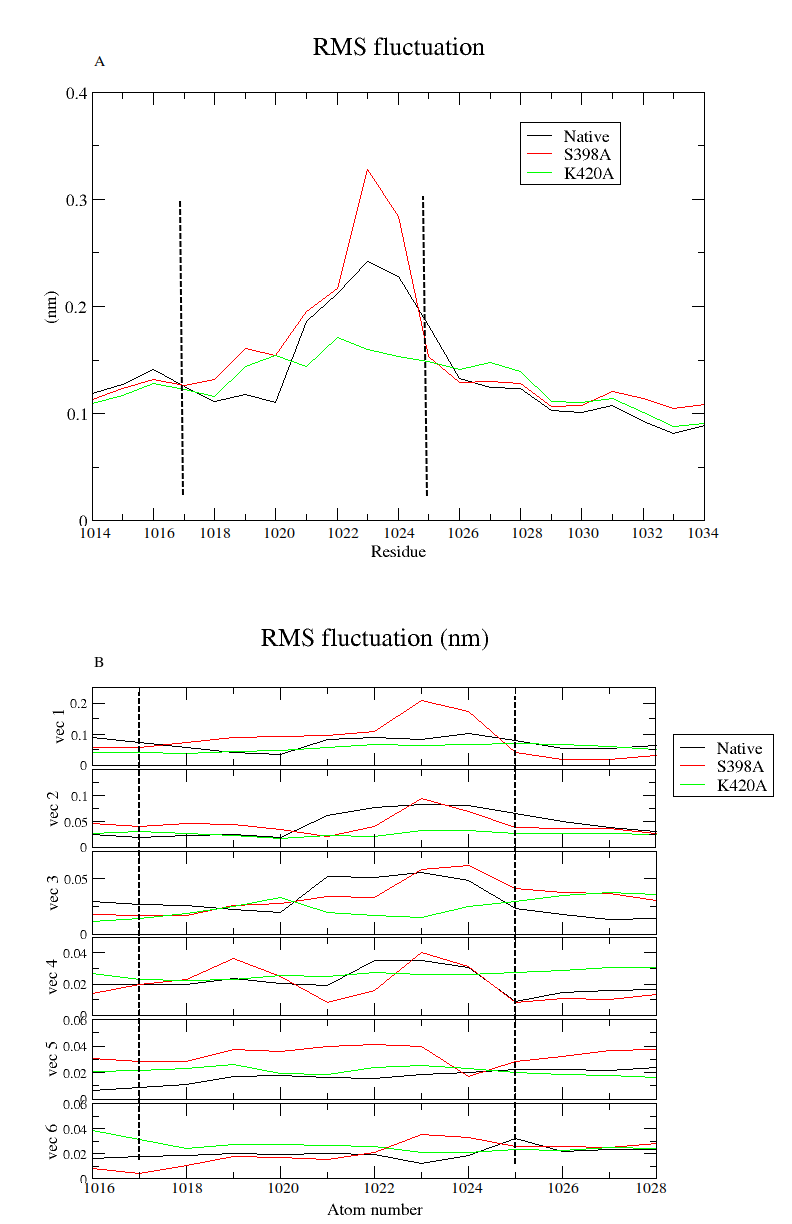

### Figure S4.png

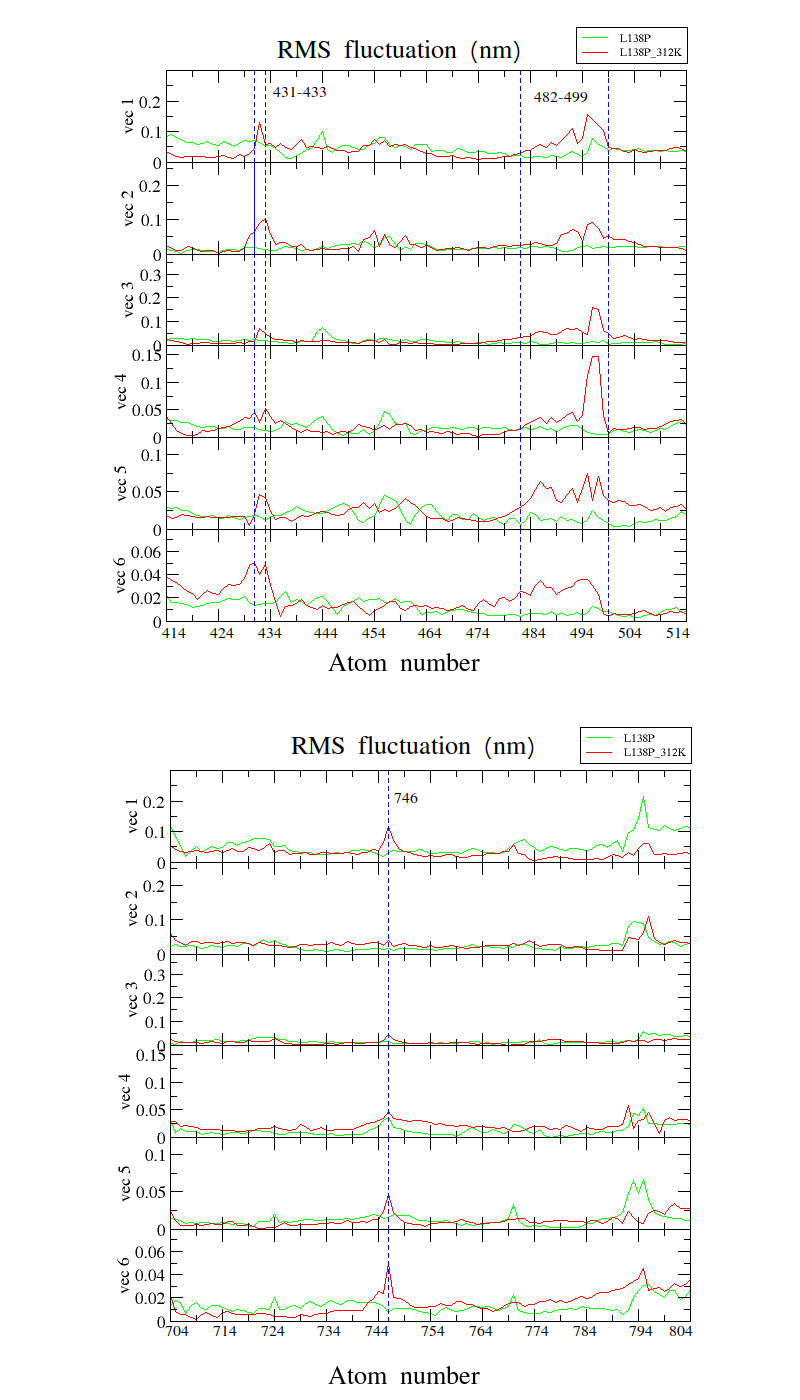

### Figure S5.pdf

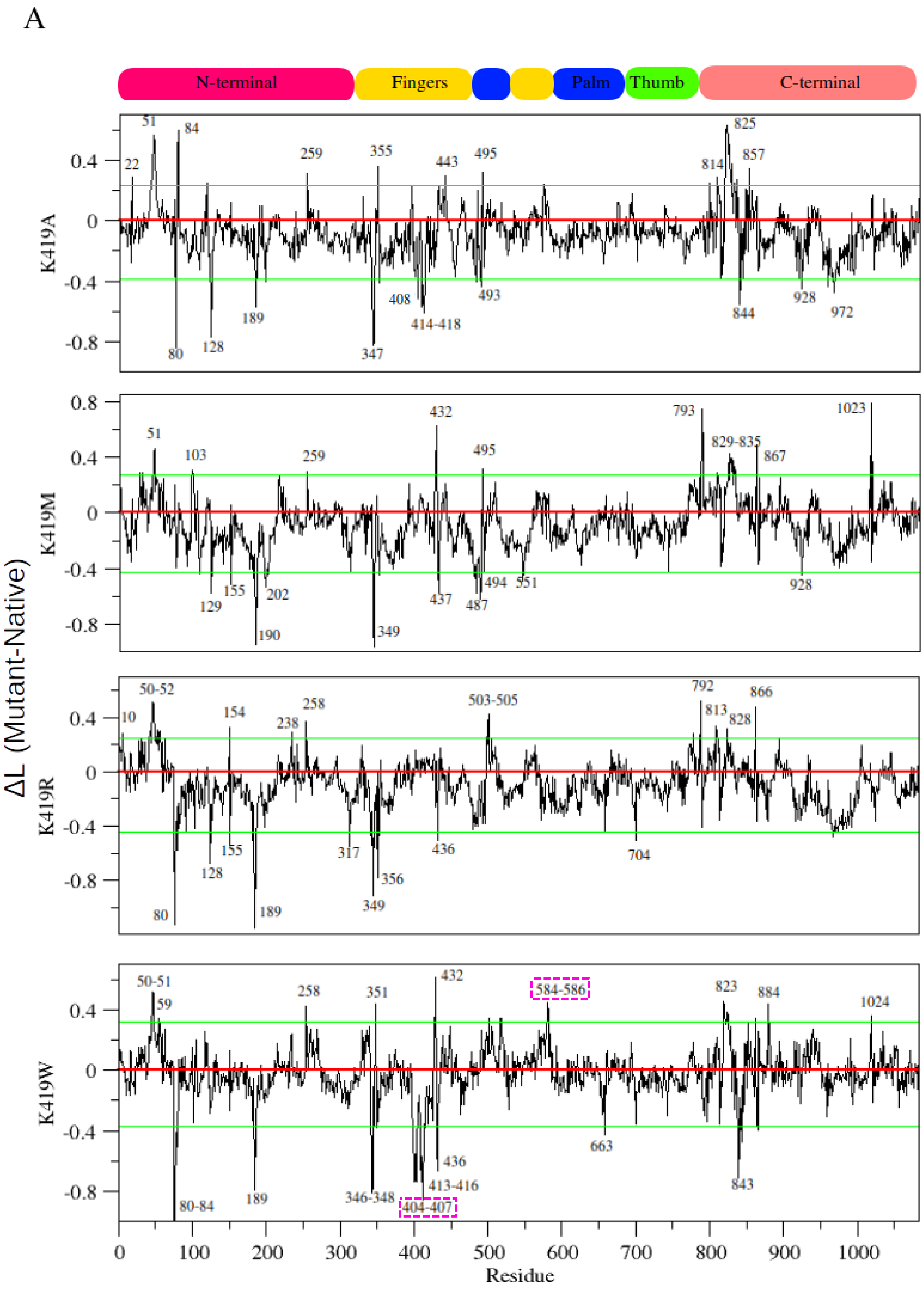

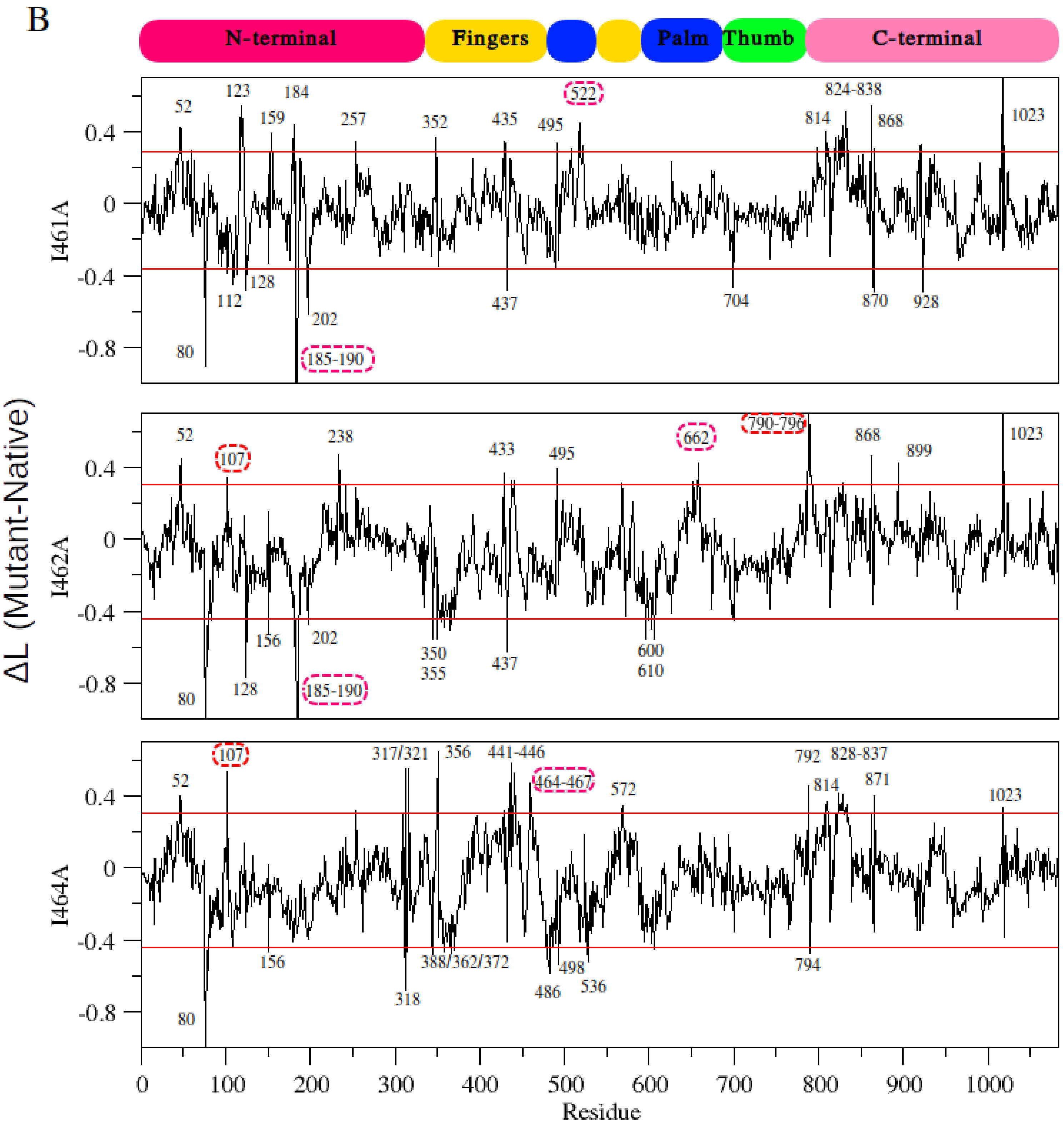

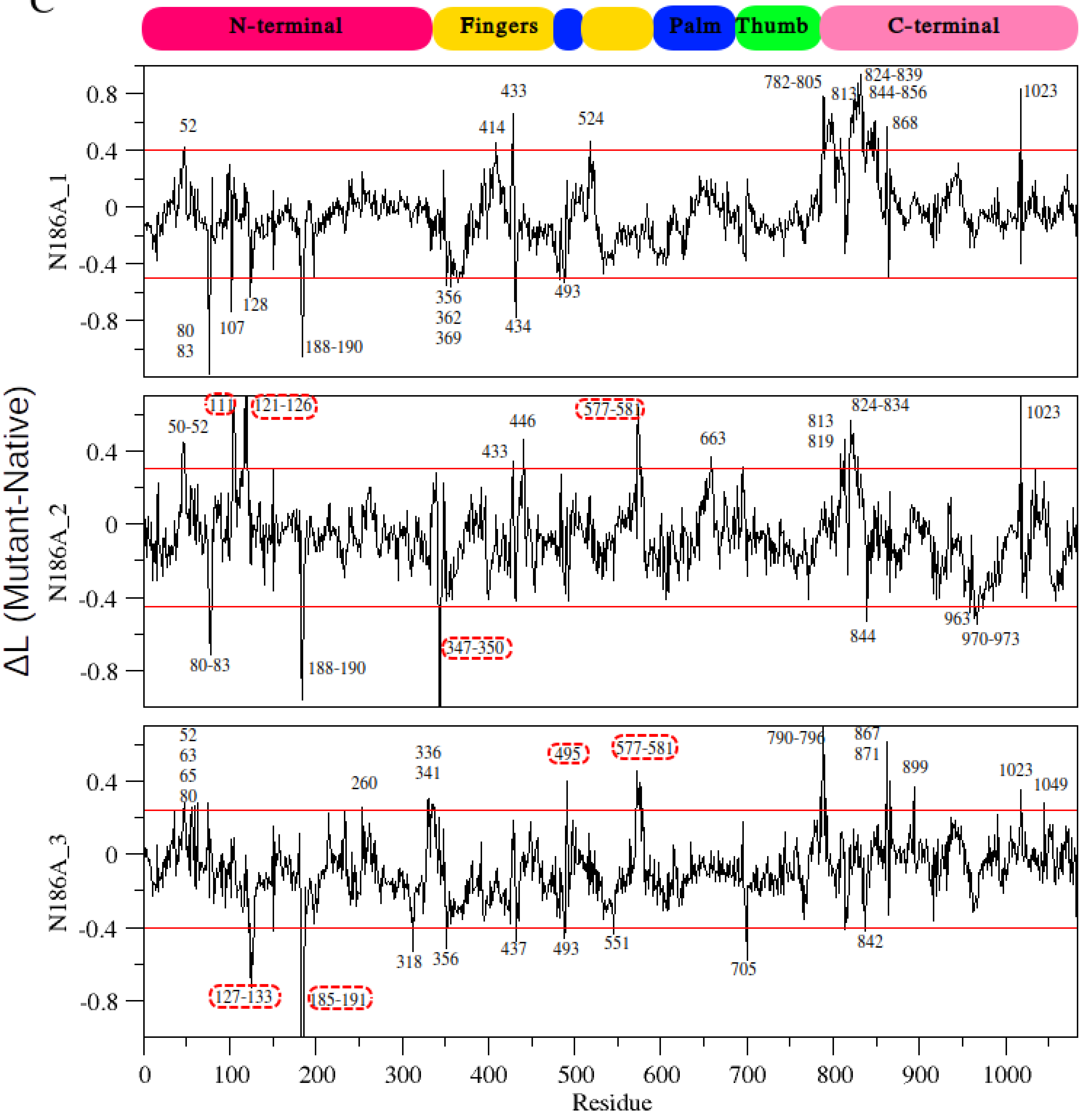

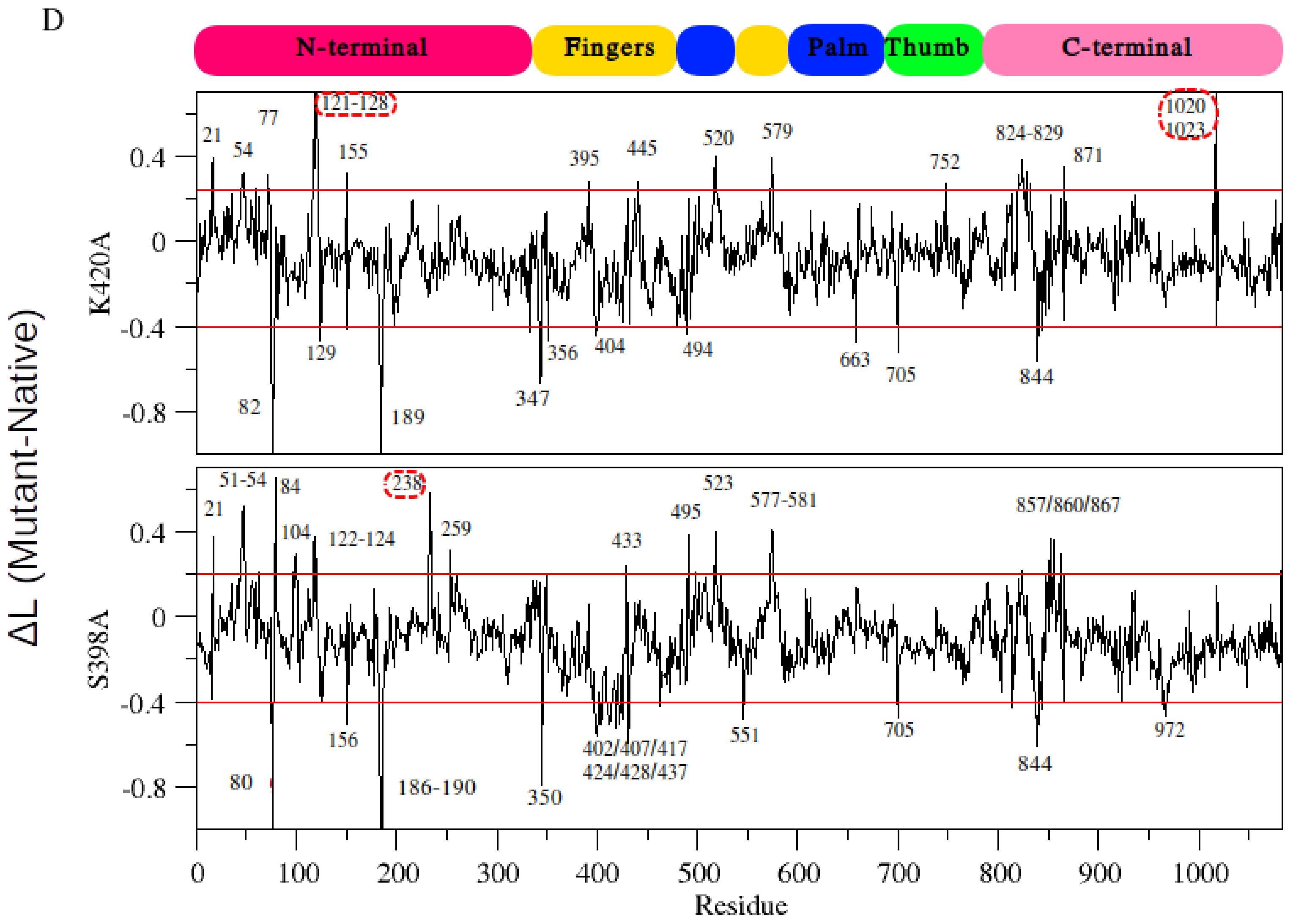

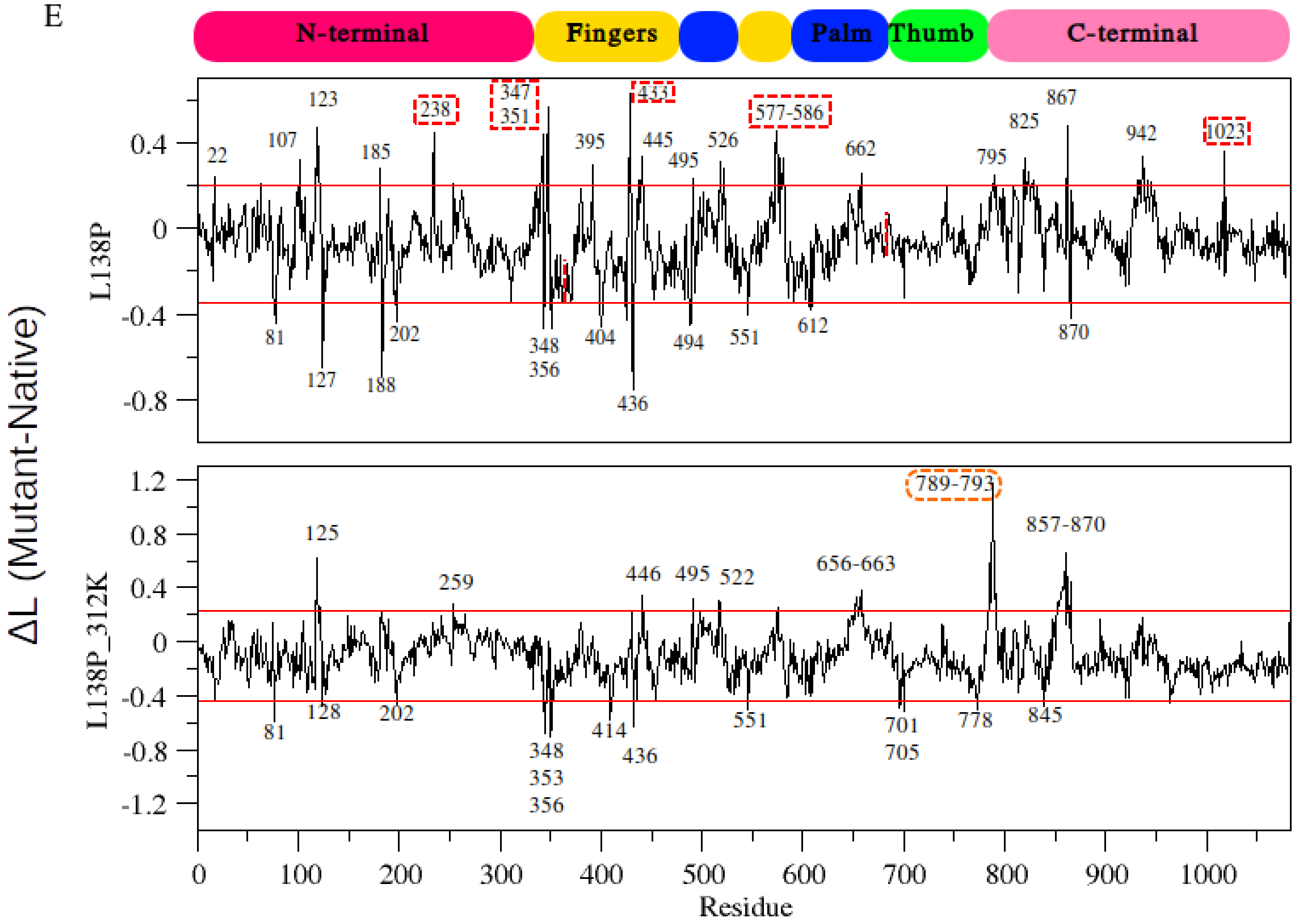

### Figure S6.pdf

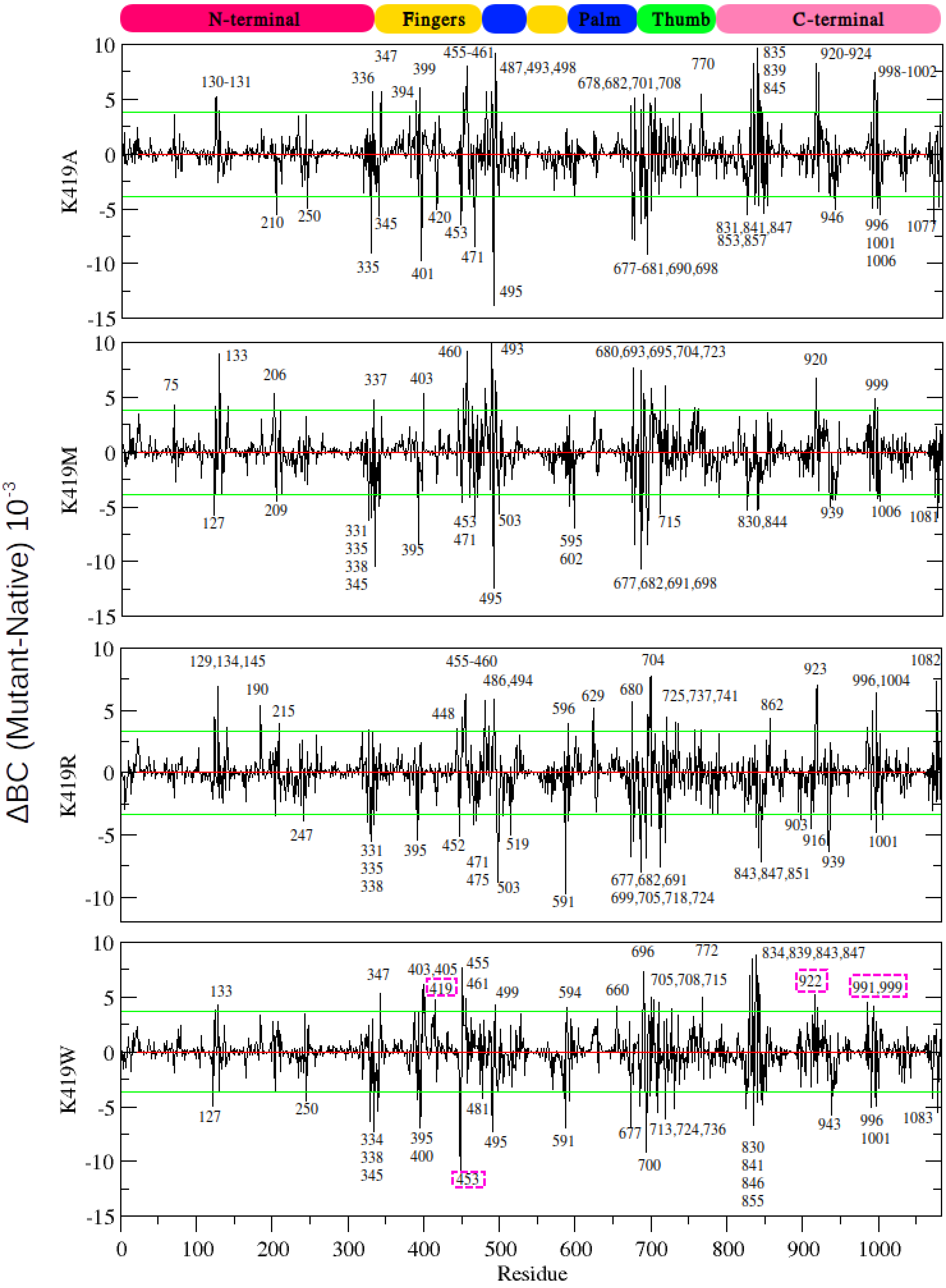

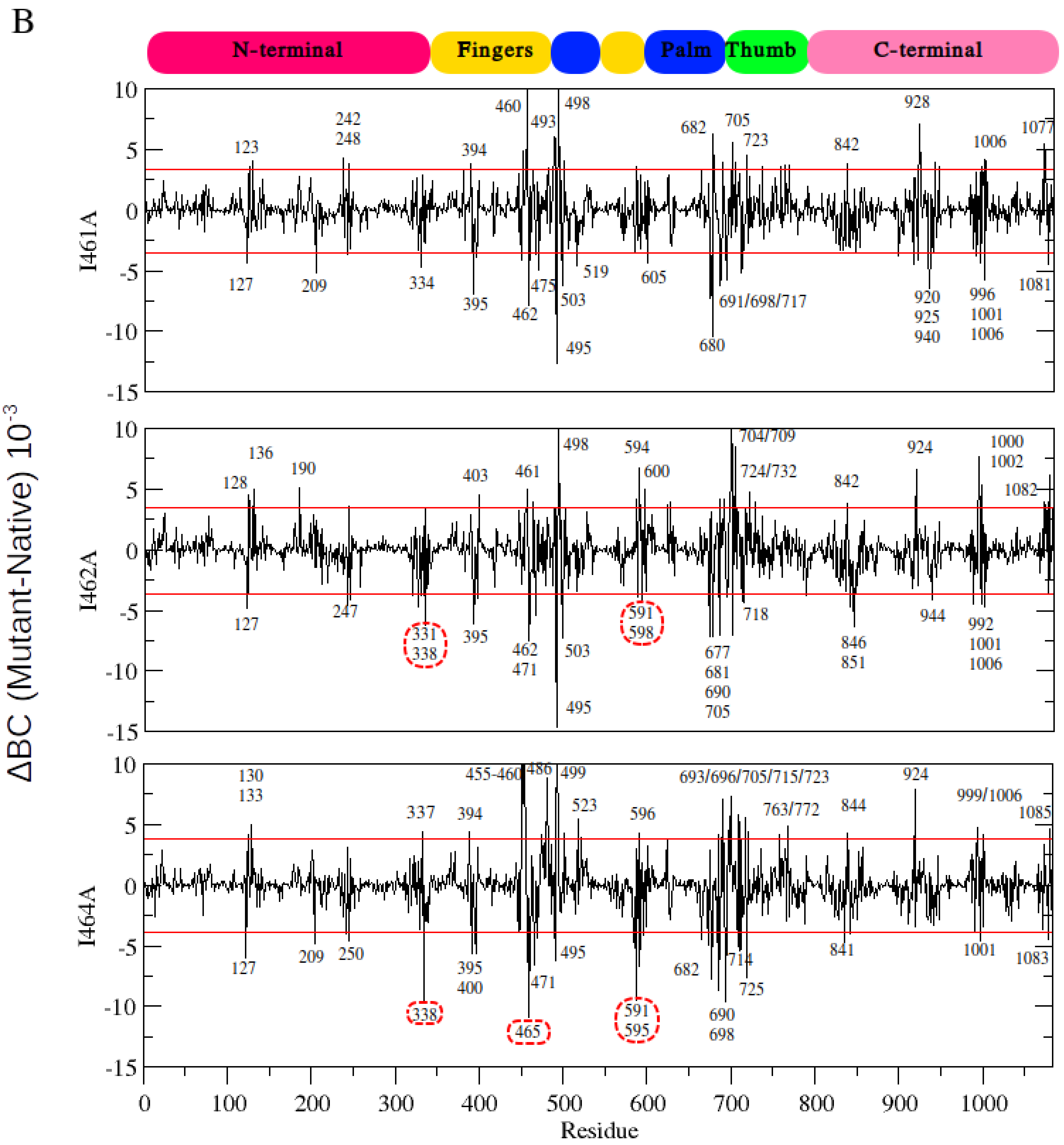

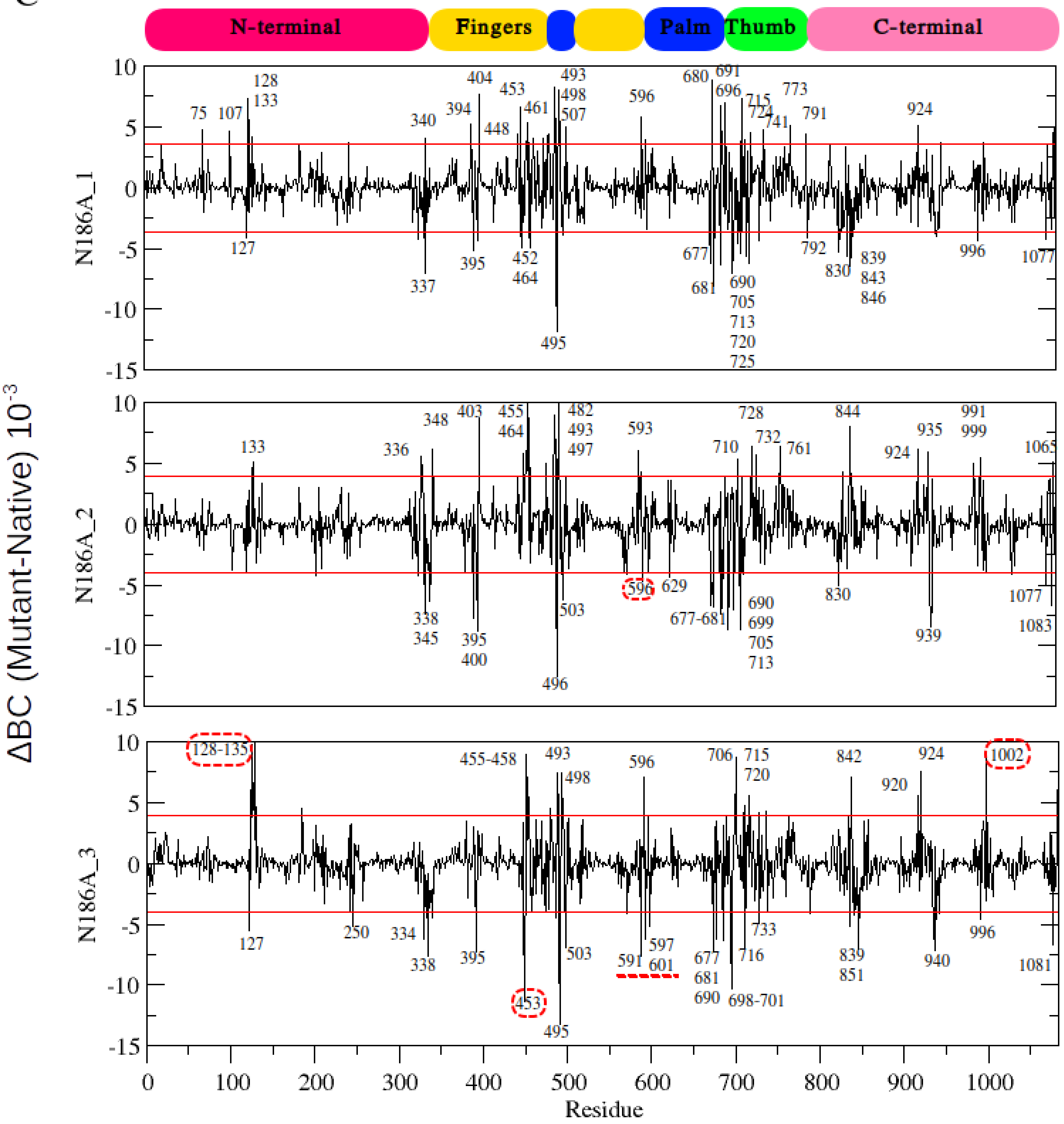

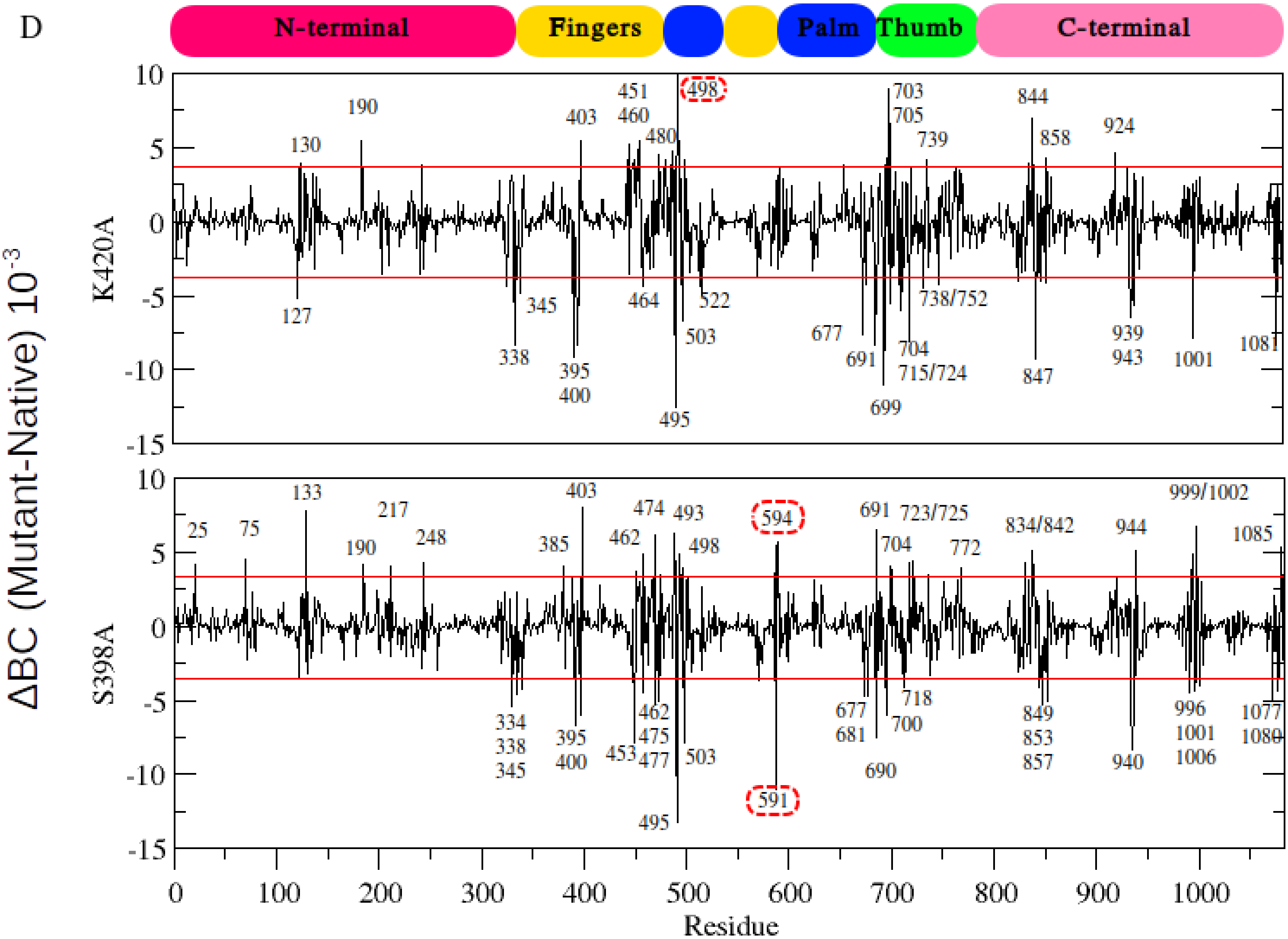

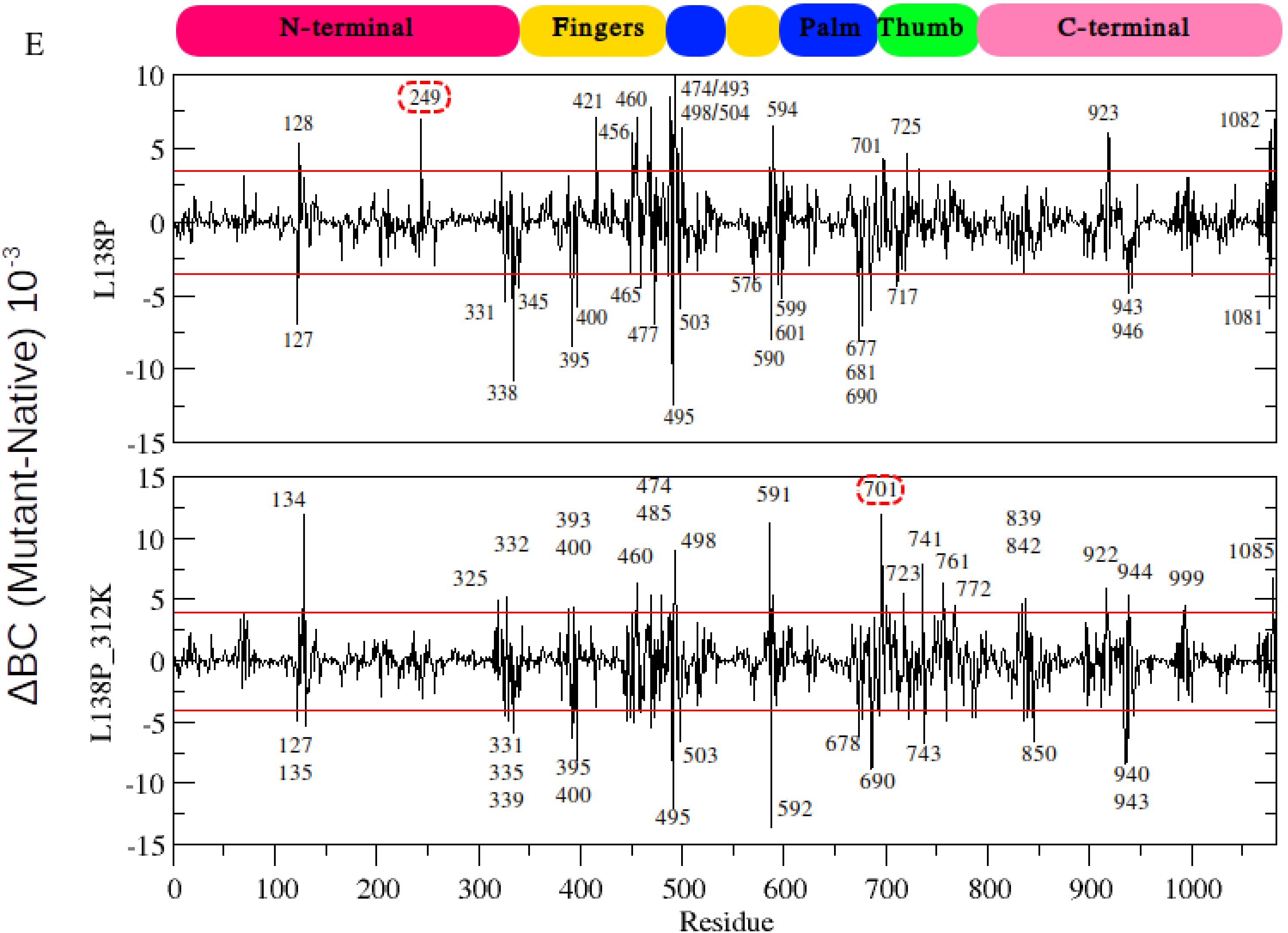

### Figure S7.pdf

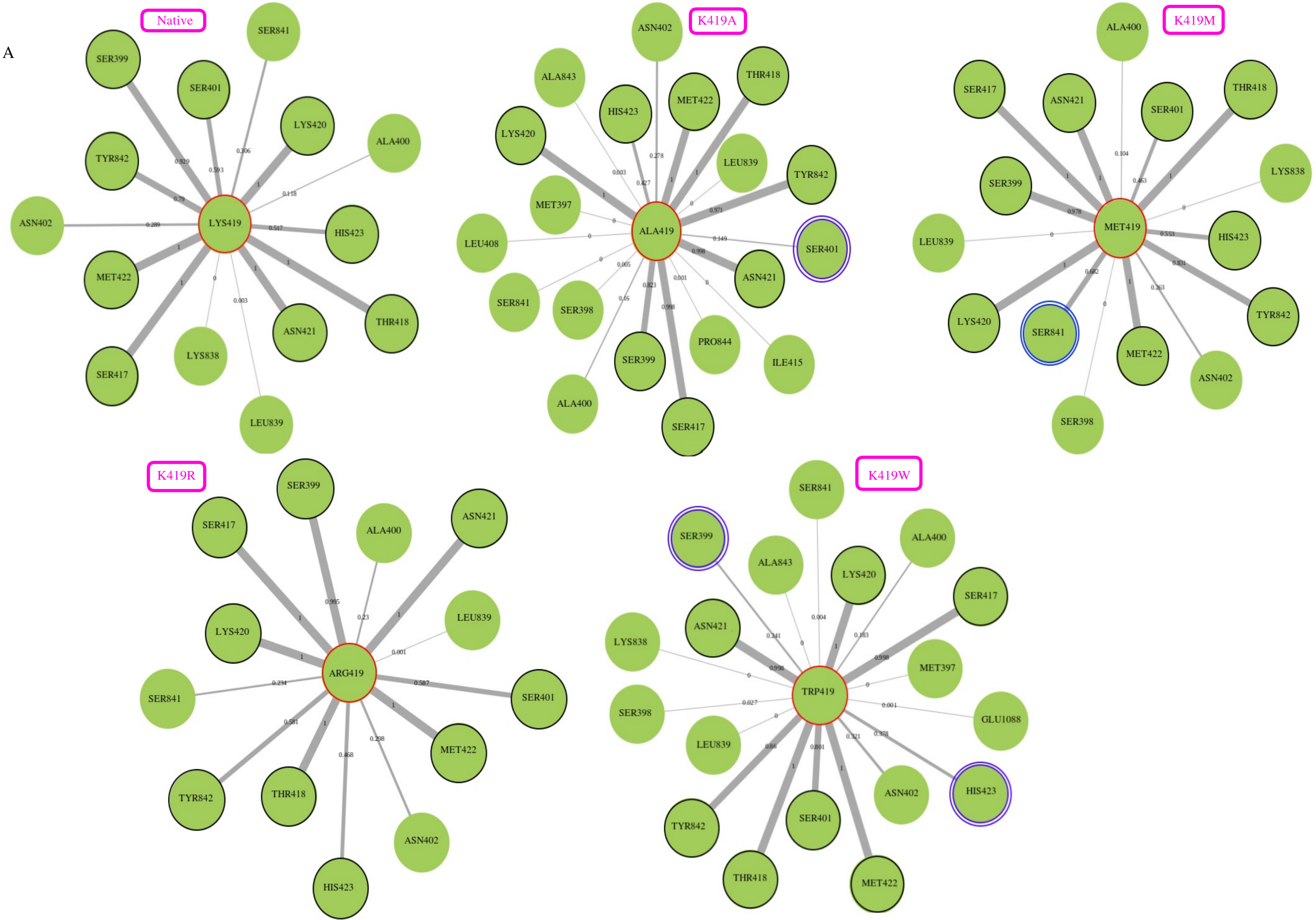

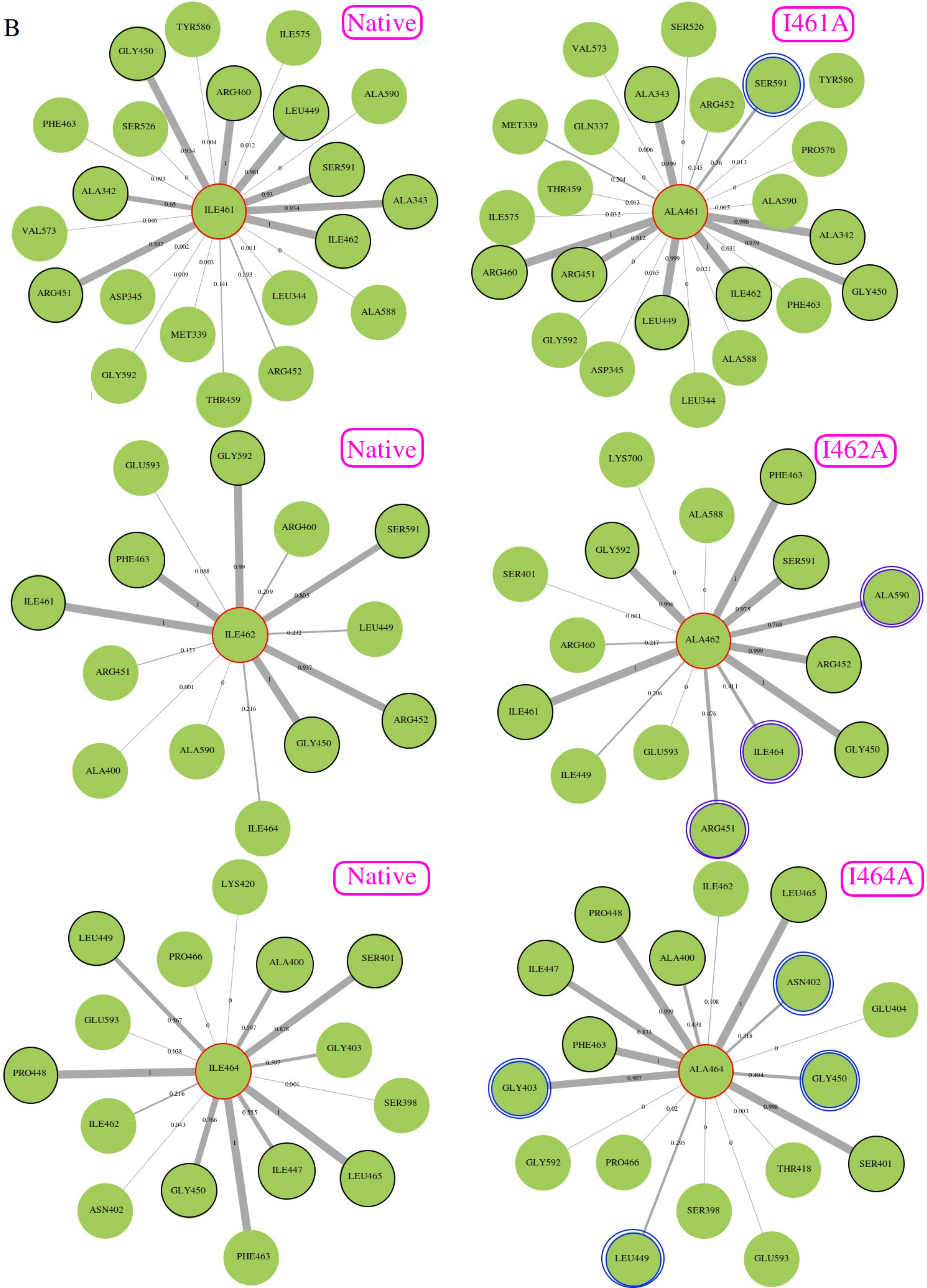

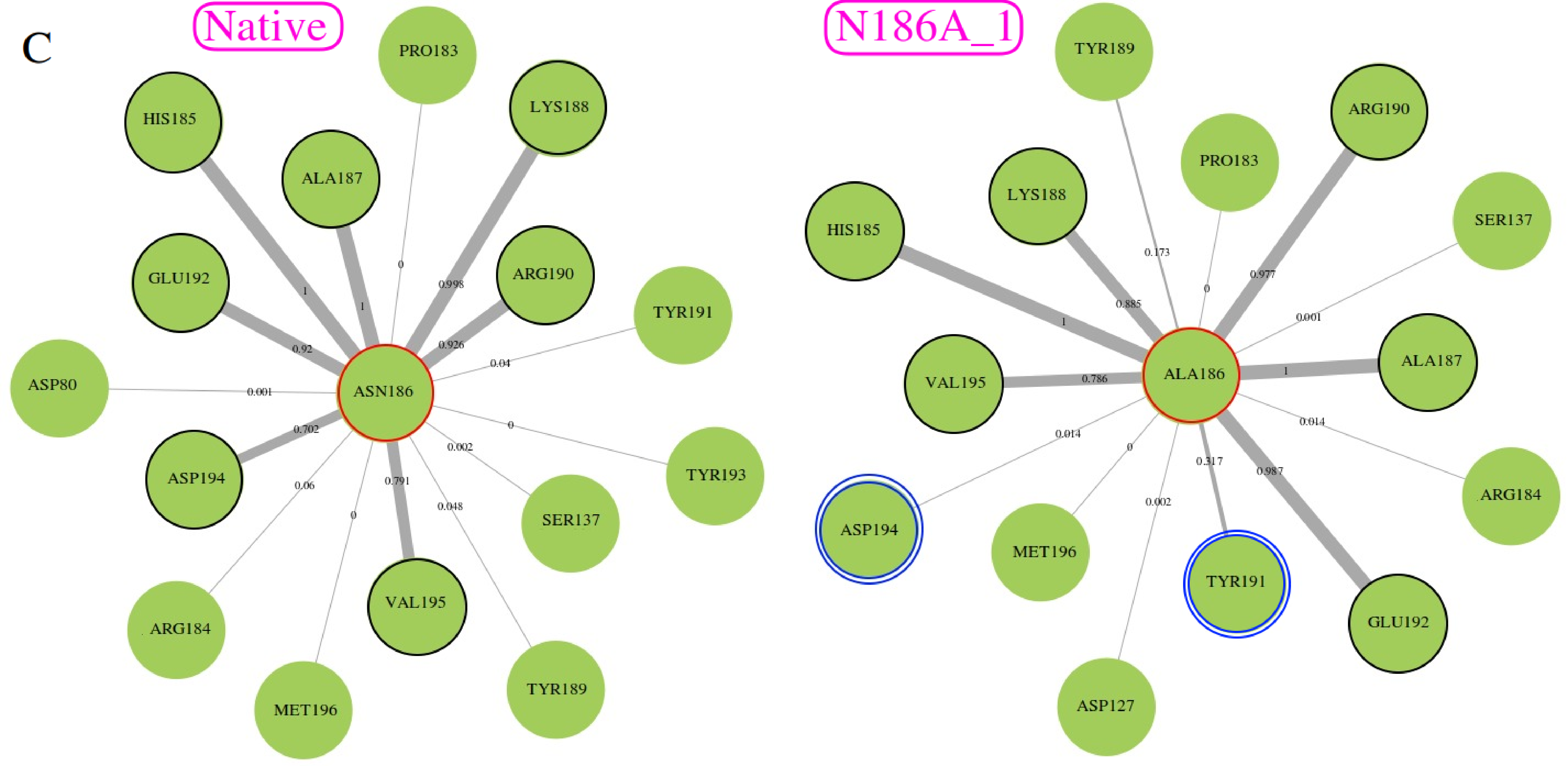

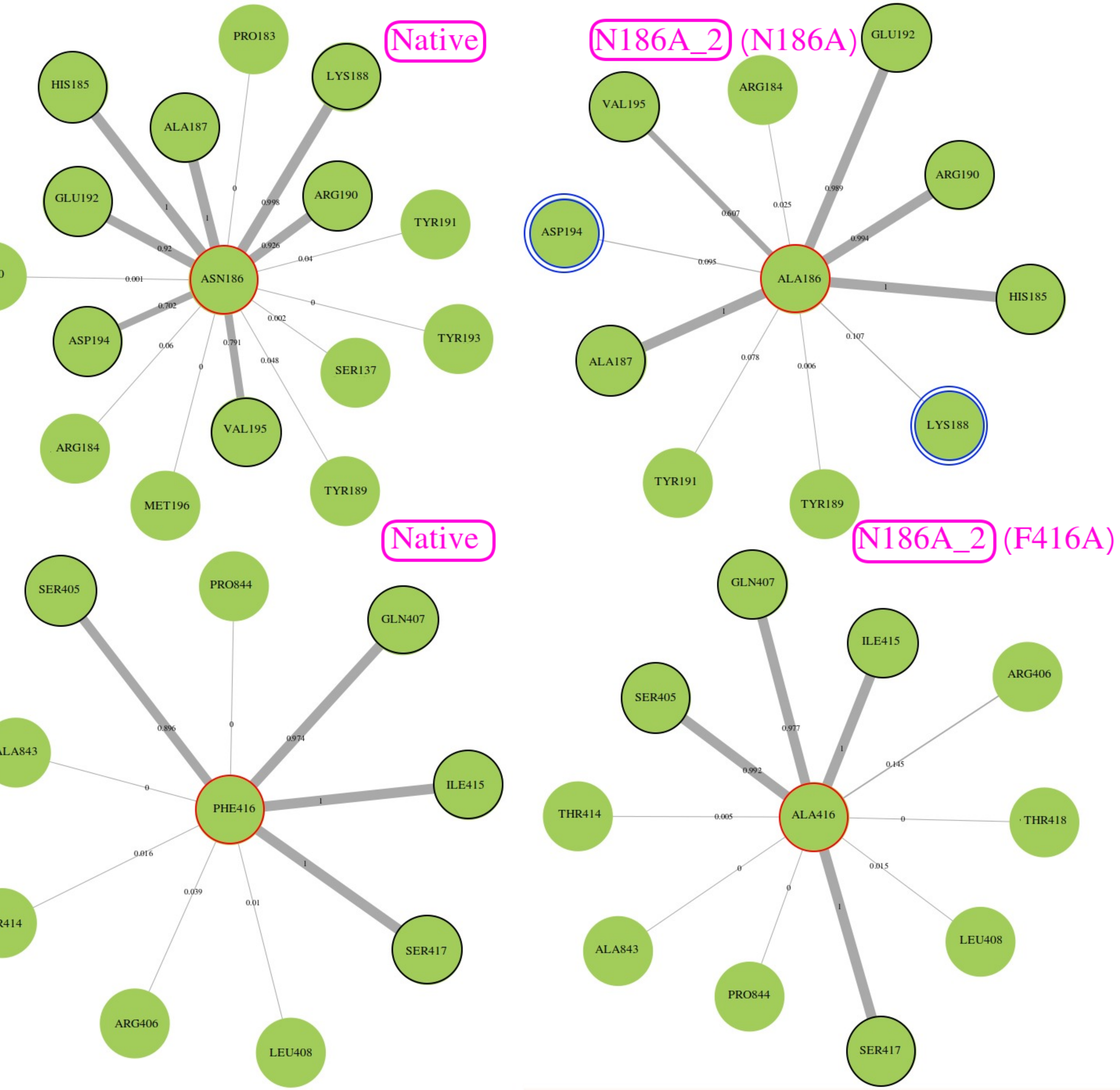

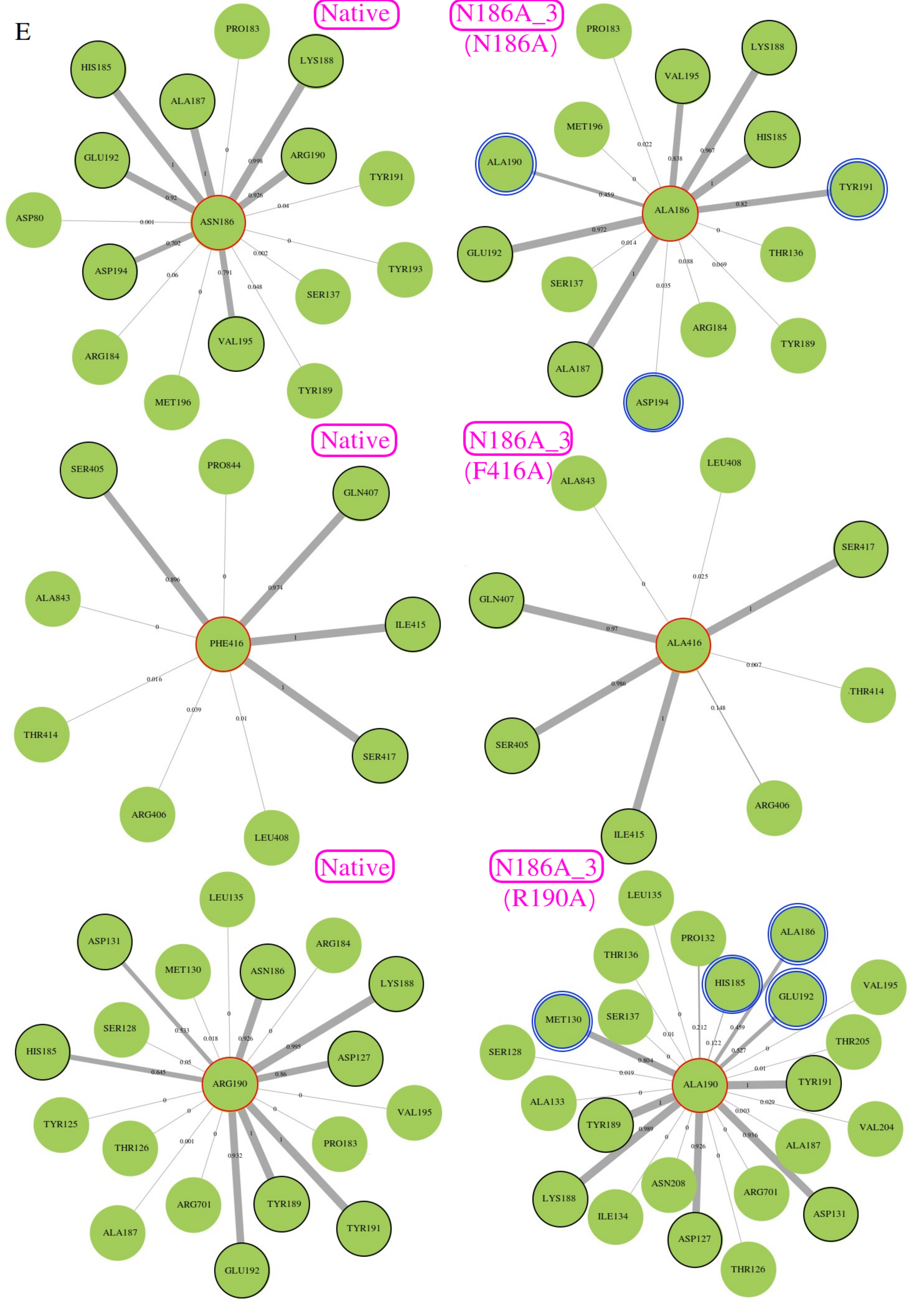

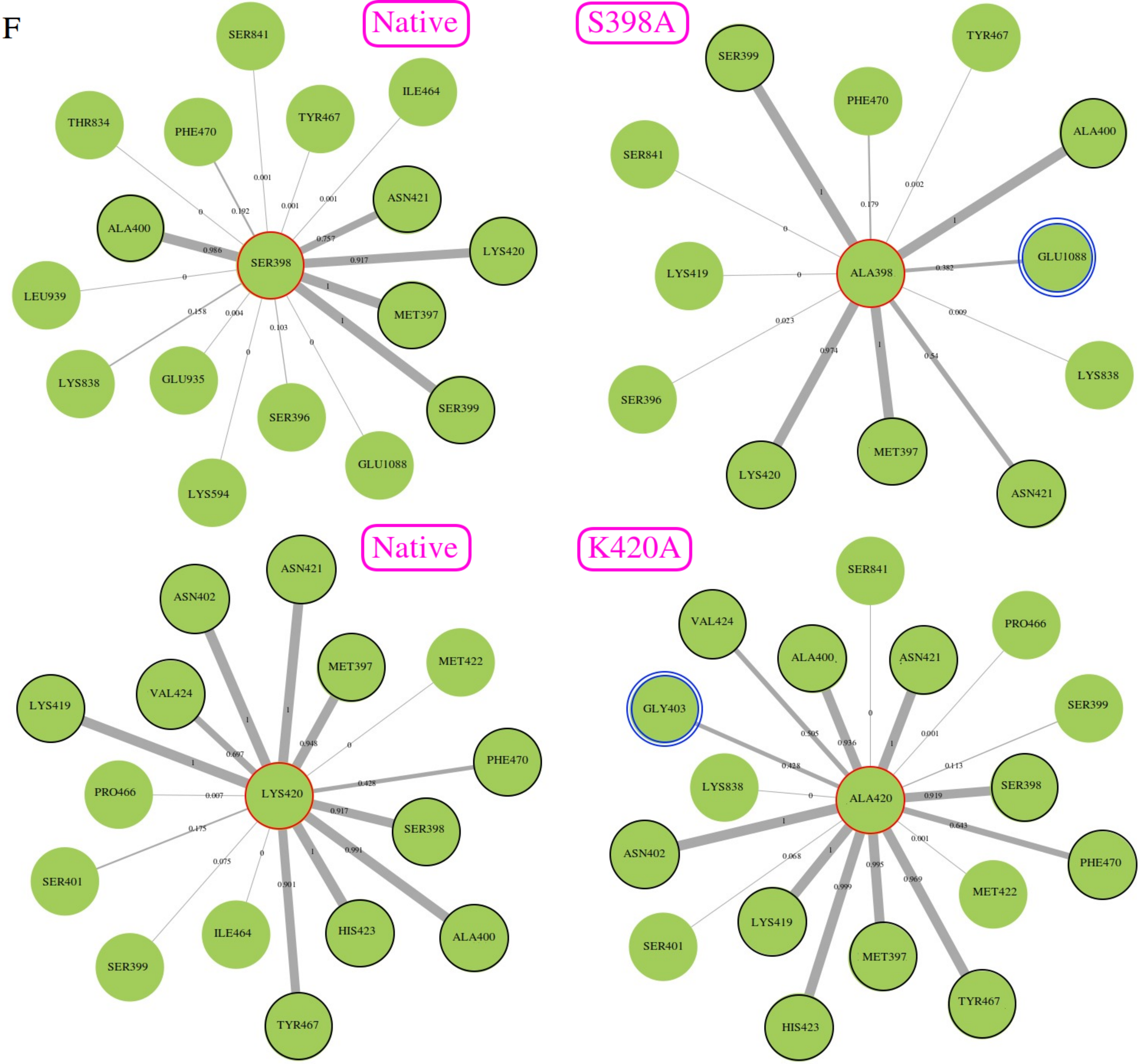

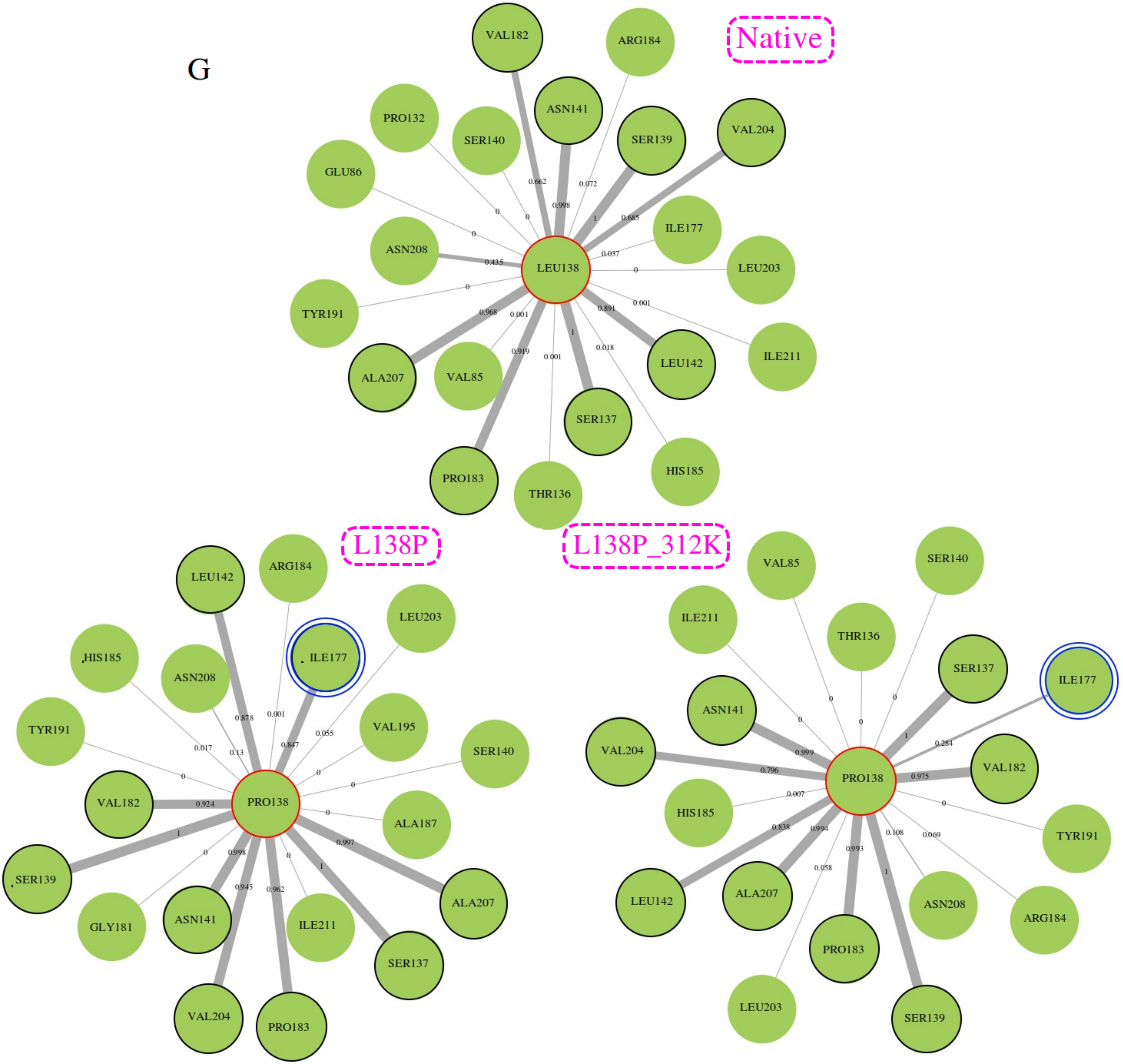

### Figure S8.pdf

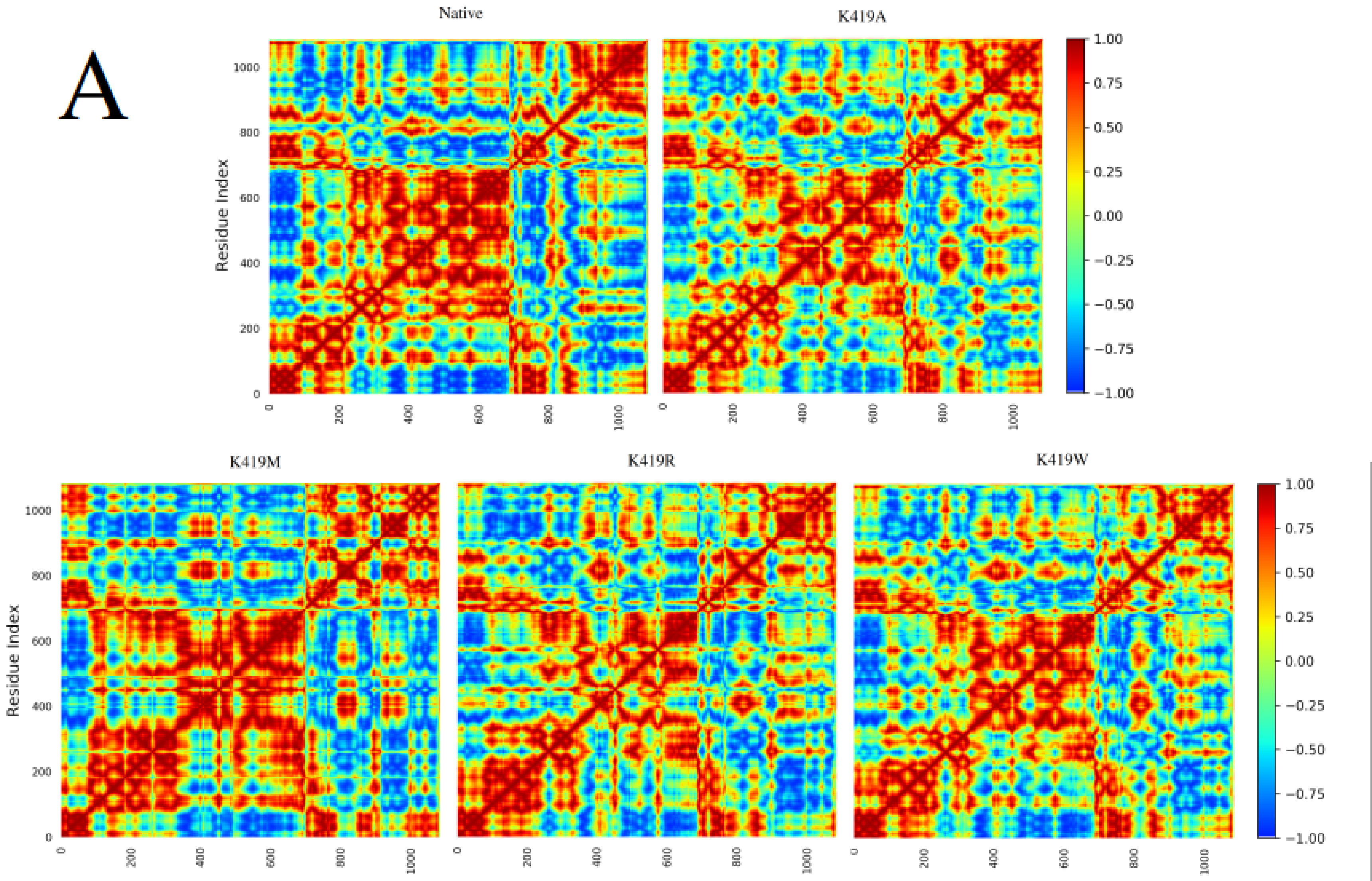

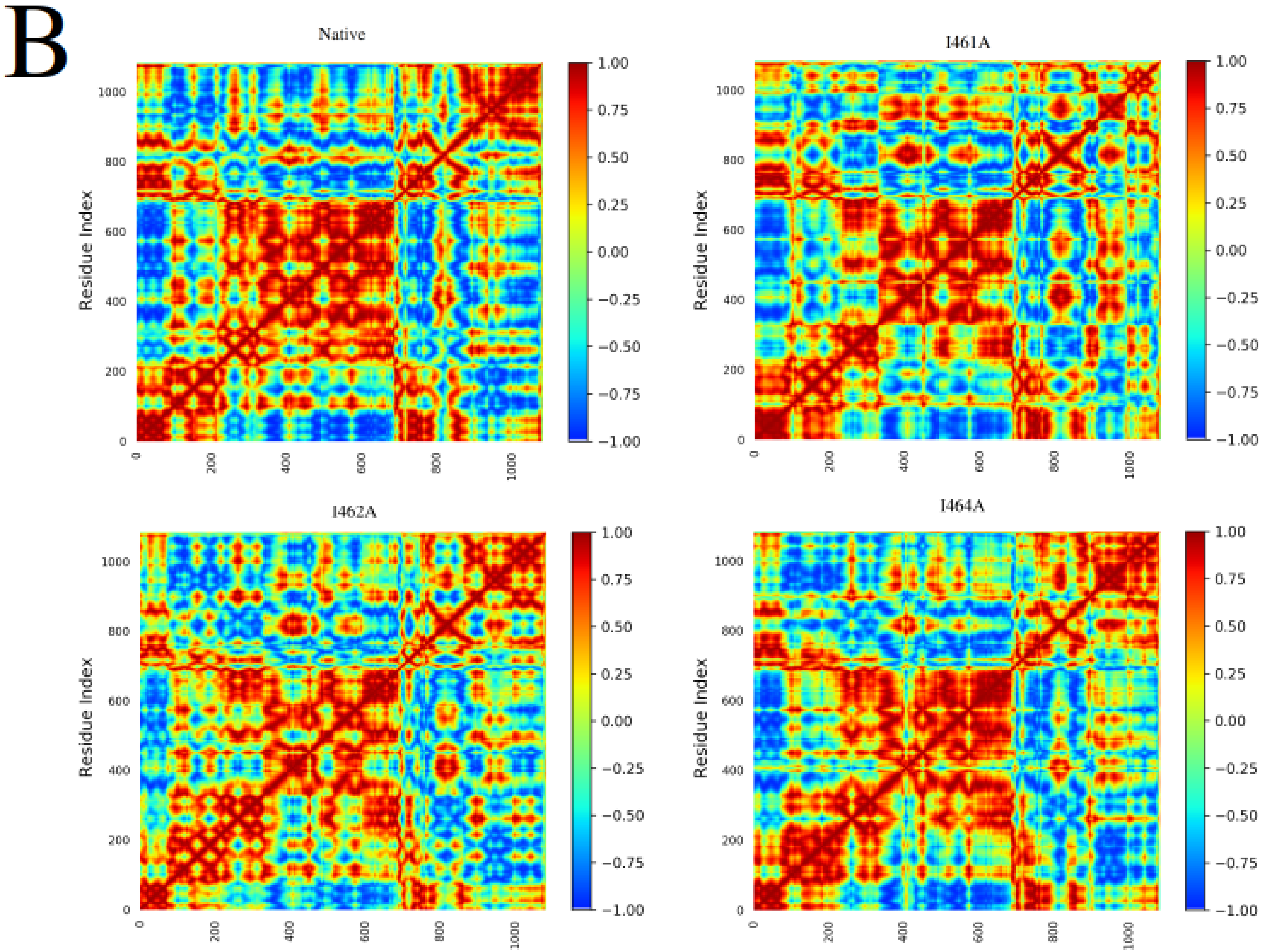

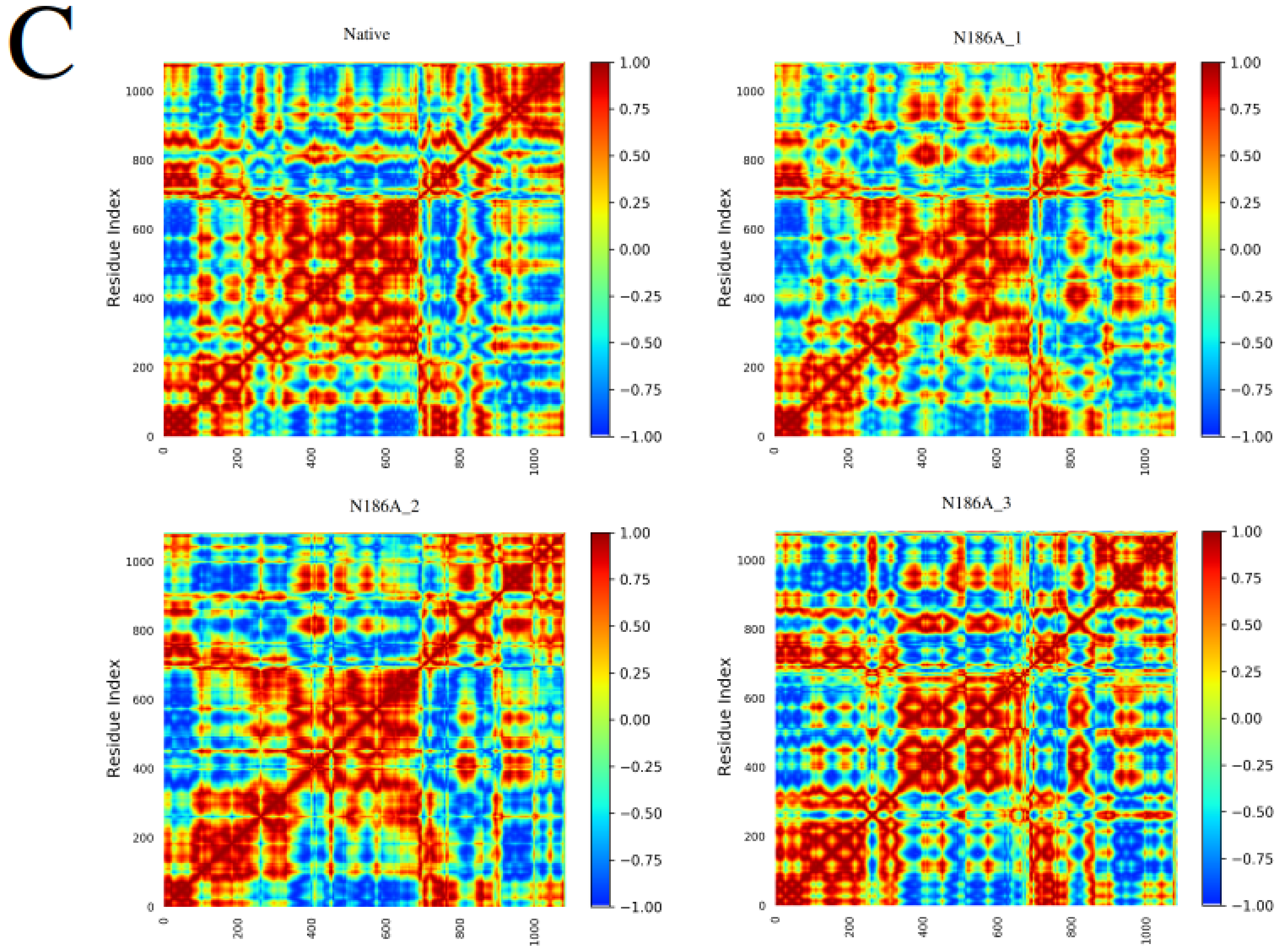

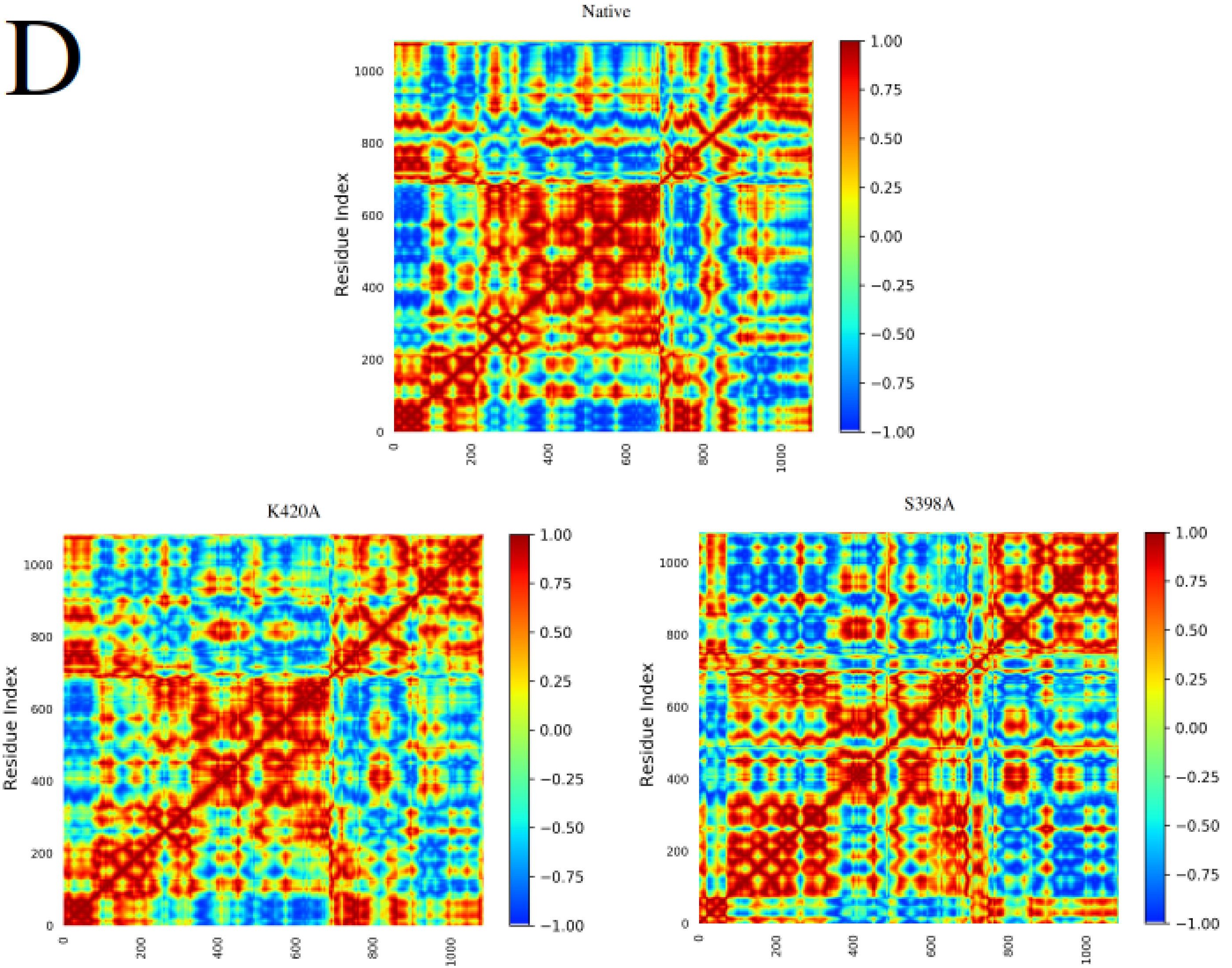

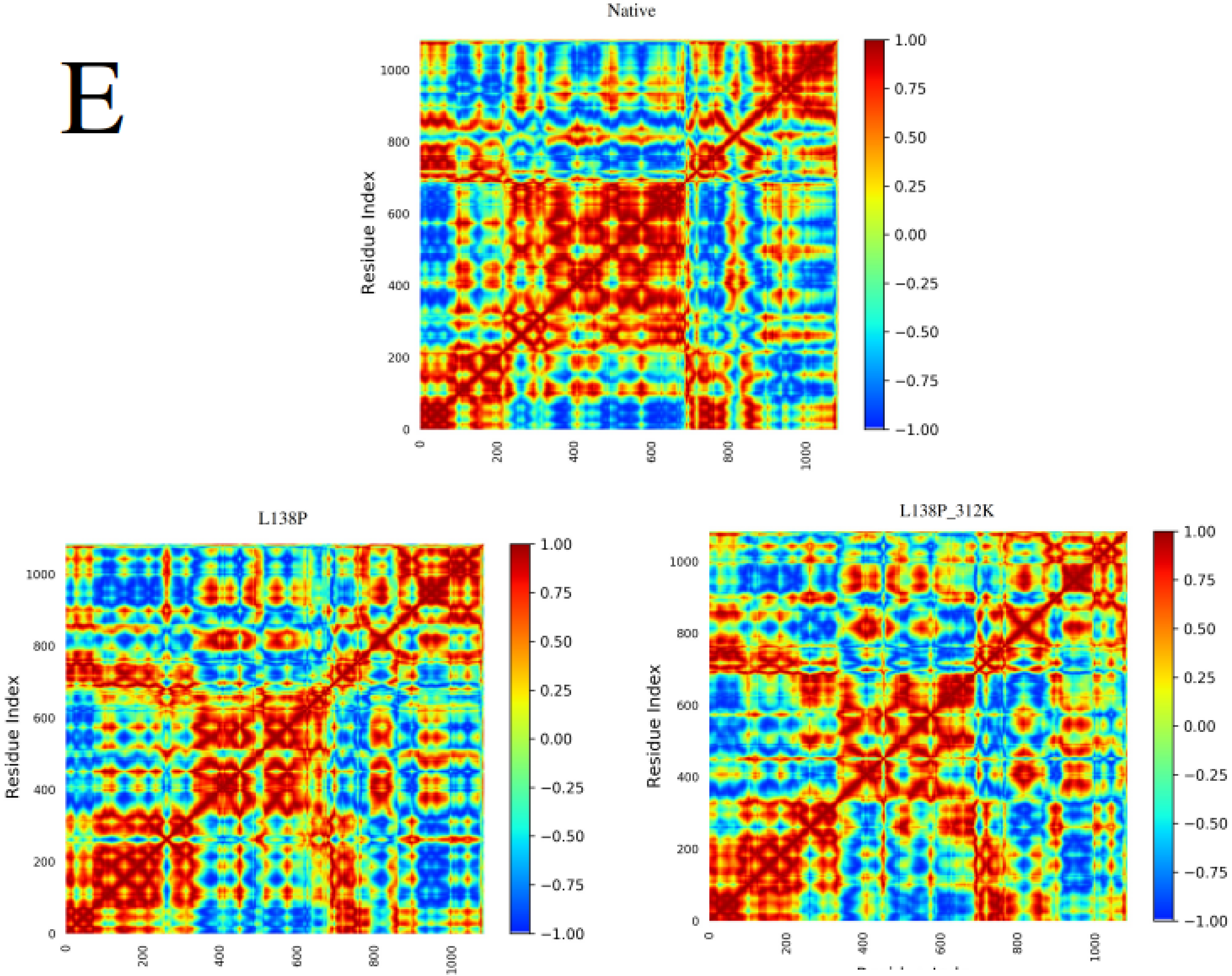

### Figure S9.png

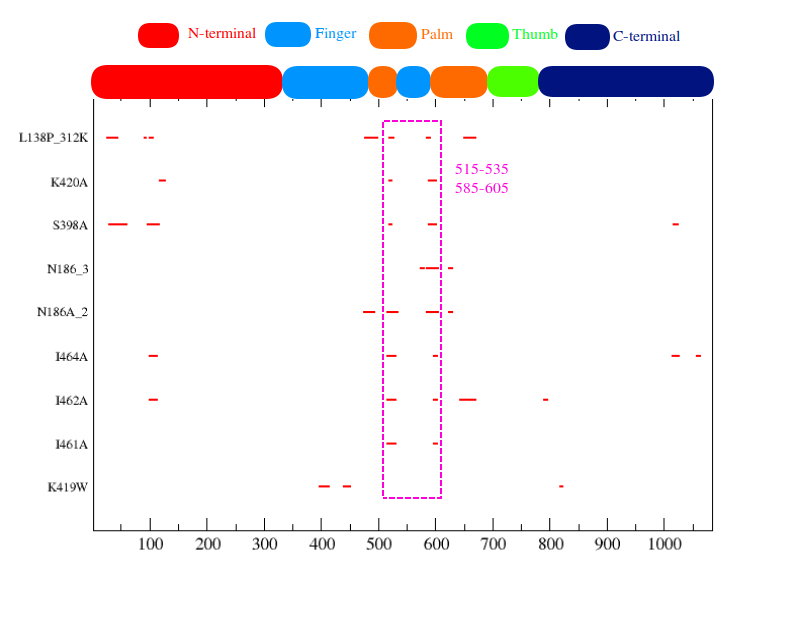
